# Connectomics by Sequencing Reveals Self-Organizing Principles of Neuronal Networks in Cerebral Organoids

**DOI:** 10.64898/2026.10.01.755792

**Authors:** Ábel Vértesy, Ramsey Najm, Bálint Doleschall, Sergey Isaev, Michael Zabolocki, Maria Novatchkova, Thomas R. Burkard, Christian Ziegler, Simone Krautberger, Chong V. Li, Igor Adameyko, Juergen A. Knoblich

## Abstract

Functional neural networks emerge as developing neurons form synaptic connections. Revealing how these connections are organized and perturbed in disease requires linking single-neuron connectivity to molecular state across thousands of networks, which is challenging with current methods. We developed COSCO (Connectomics by Sequencing in Cerebral Organoids), combining barcoded monosynaptic rabies tracing with single-cell RNA sequencing to reconstruct over 10,000 networks across organoids. We demonstrate that organoid circuitry self-organizes into stereotyped network motifs governed by cell-intrinsic wiring rules aligning with features of mouse cortical connectivity. Projecting network topology onto transcriptomic space reveals that wiring correlates with cell-adhesion and synaptic gene expression programs. In a Tuberous Sclerosis Complex model, pathological hyperexcitability coincides with topological rewiring: mutant networks become more heterogeneous, driven by the transcriptional distribution of upper layer excitatory cell states. This work establishes cerebral organoids as a model to study self-organization of human connectivity, and its disruption in neurodevelopmental disease.

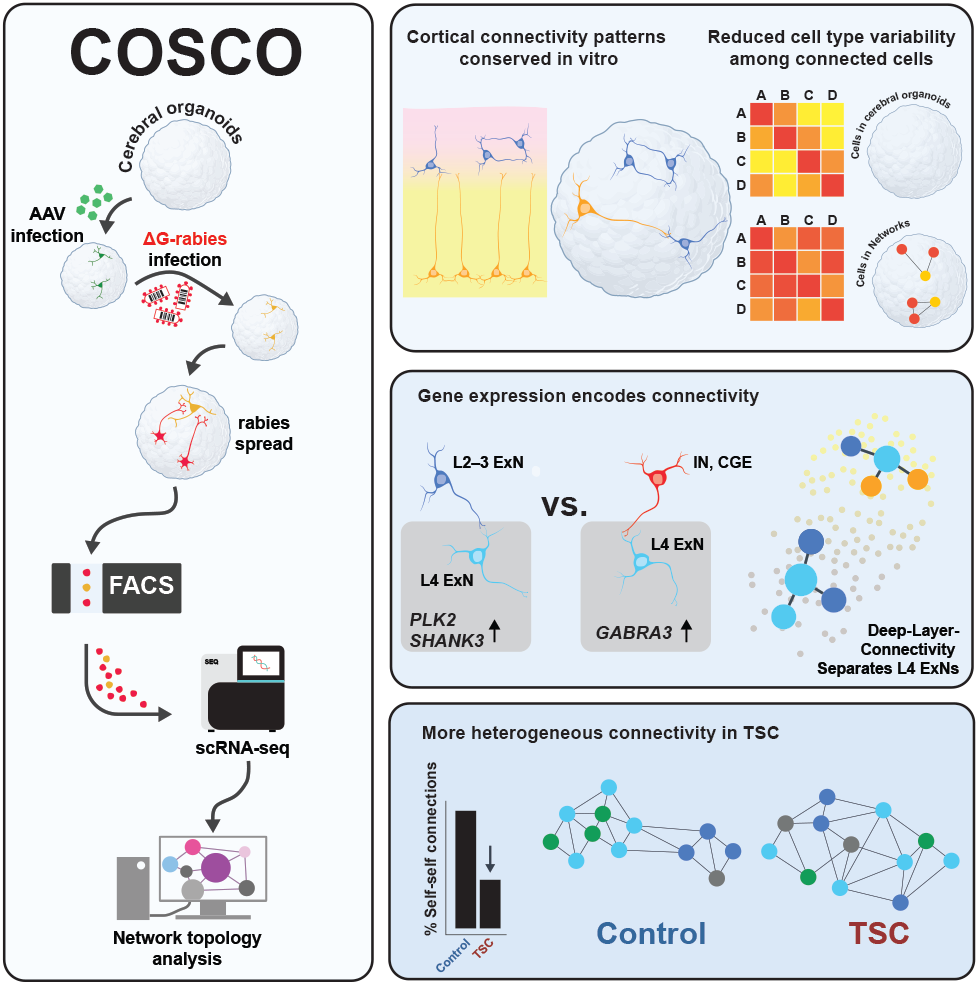

## Introduction

The human brain’s capacity for cognition and behavior emerges from the organization of billions of neurons into an interconnected network known as the connectome ^1^. The assembly of these networks in mammals is not completely hardwired, but probabilistic, requiring a substantial degree of self-organization that is vulnerable to perturbation ^2–5^. Understanding the mechanism underlying this self-organizing process in humans depends on two key requirements. One is a human-specific system that permits network formation in the absence of external stimulation. The other is large-scale, joint measurement of connectivity and transcriptomic state at single-cell resolution. Despite its importance, the assembly of human neural networks is poorly understood as both criteria remain unmet: the human brain is difficult to access and develops with external cues, while existing methods do not resolve connectivity and molecular identity at scale.

Cerebral organoids derived from human induced pluripotent and embryonic stem cells (hiPSCs, hESCs) provide a tractable system to study neuronal self-organization directly ^6–8^. They recapitulate key features of cortical development ^9^, including the maturation of diverse cell types ^10^, migration ^11–13^, and complex spontaneous network activity ^14,15^. Moreover, they are an effective tool to model neurodevelopmental diseases ^16,17^. For example, organoids derived from patients with Tuberous Sclerosis Complex (TSC), an mTORopathy characterized by epilepsy and neurodevelopmental impairment, recapitulate key morphological and electrophysiological disease phenotypes ^18–21^, suggesting underlying alterations in network connectivity. As organoids develop in the absence of external stimulation, and can model functional pathology in neurodevelopmental disease, they are a powerful system for revealing intrinsic principles of network assembly and their disruption.

There has been significant progress in understanding the molecular mechanisms of synaptogenesis ^22,23^, and in describing connectivity in precise spatial detail by electron ^24,25^ and expansion microscopy ^26,27^. Yet joint readout of connectivity and transcriptome at scale require sequencing-based connectomics ^28^. Scalable single-neuron projectome mapping is well established ^29–31^ but mapping synaptic connectivity between interconnected neurons relies primarily on the monosynaptic retrograde tracing (MRT) system, which leverages glycoprotein-deficient rabies virus (RVΔG) ^32,33^. By labeling rabies virions with a unique barcode and recovering those using single-cell RNA sequencing (scRNA-seq), one can identify groups of cells with identical barcodes, and thus reconstruct synaptic networks of individual human neurons ^34^. Recent approaches have recovered hundreds of presynaptic networks of single postsynaptic neurons ^34–37^. To increase scale and map thousands of networks, we developed COSCO (**Co**nnectomics by **S**equencing in **C**erebral **O**rganoids). Therein, we combined and innovated on sparse labeling, experimental-, viral-, barcode design, and introduced novel computational methods, including latent network embedding by clone2vec ^38^. With this approach, we reconstructed over 10,000 unique neuronal networks across organoids and cell lines and demonstrated that aspects of connectivity patterns *in vivo* are recapitulated *in vitro*. With clone2vec, we revealed that networks in organoids segregate into recurrent patterns of connectivity which is embedded in the transcriptome. In the context of TSC, COSCO uncovered a shift toward less constrained wiring driven by changes in network cell-state composition. This network reorganization was predictive of observed electrophysiological abnormalities. Together, COSCO maps thousands of neuronal networks with single-cell transcriptional resolution in a human-specific model system, enabling one to dissect the mechanisms of self-organization, and its perturbation in disease.

## Results

### Establishing connectomics-by-sequencing in cerebral organoids

MRT maps direct presynaptic inputs using an EnvA-pseudotyped, glycoprotein-deleted rabies tracer virus carrying a fluorophore (e.g., RVΔG–E2-Crimson) and an adeno-associated virus (AAV) helper. Because the glycoprotein (N2cG) is deleted, RVΔG cannot spread autonomously. RVΔG is then pseudotyped with EnvA, an avian virus’s envelope protein, absent from mammalian cells. Consequently, it can only infect cells expressing its cognate receptor, TVA, delivered by the AAV. The helper therefore generates “putative starter cells” by expressing TVA, N2cG, and EGFP which enables RVΔG entry, monosynaptic spread, and marks helper-infected cells respectively. Helper-positive cells also infected by RVΔG become “starter cells” or “starters”, from which rabies spreads retrogradely to direct presynaptic partners regardless of dendritic synapse localization or starter-cell identity ^39^. In our experiments, RVΔG infection was performed 14 days after AAV helper delivery, followed by 7 days of monosynaptic tracing. Barcoded MRT follows this strategy but with the addition of barcodes, which we call unique viral identifiers (UVIs), integrated into the RVΔG genome. This labels individual starter cells and their direct presynaptic partners. Cells sharing a UVI can therefore be assigned to a single network **(Figure 1A)**.

**Fig. 1.**
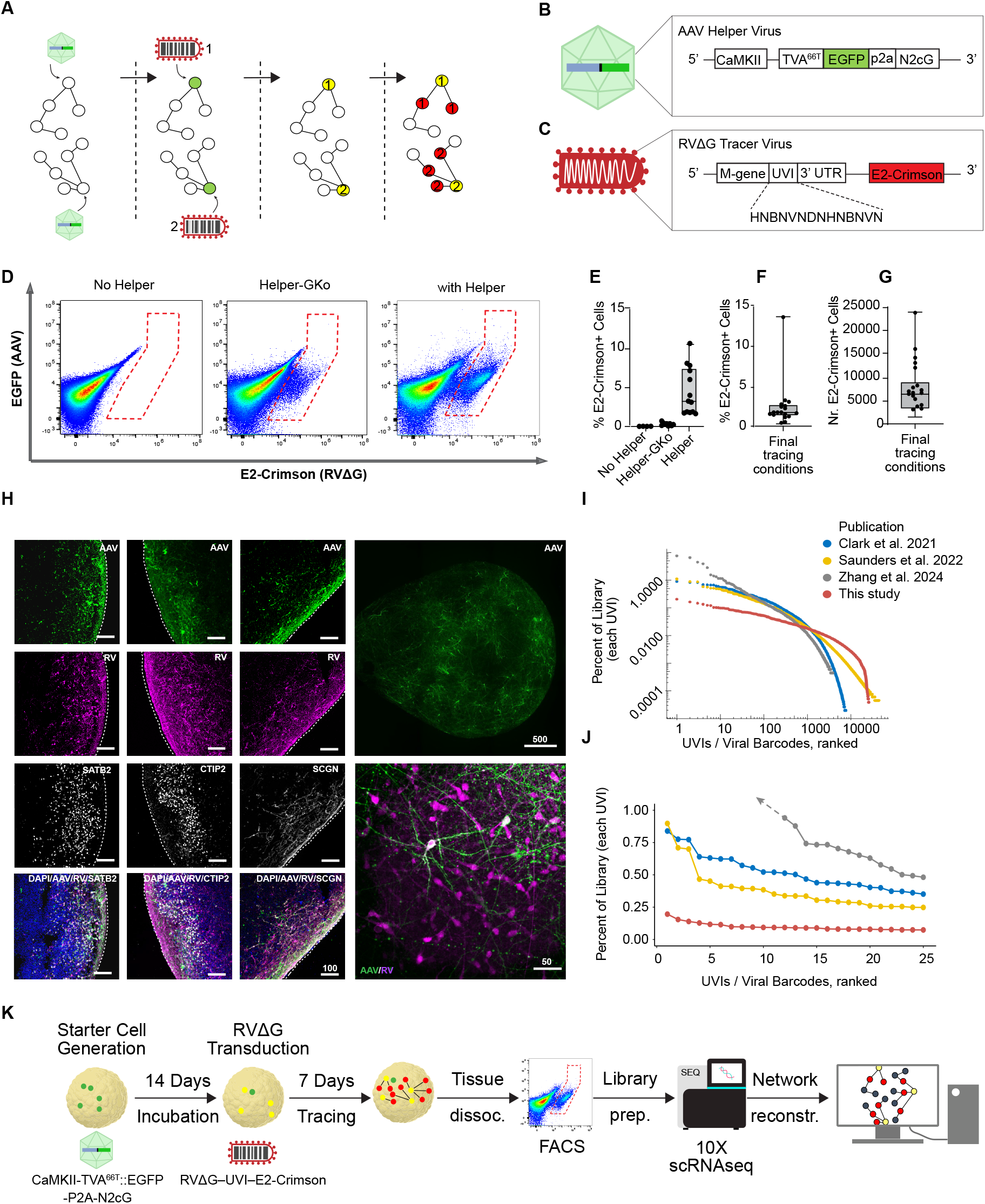
COSCO: barcoded monosynaptic retrograde tracing in cerebral organoids. **(A)** Schematic of RVΔG-based MRT. The CaMKII-driven AAVrg construct restricts putative starter cells to excitatory neurons (green); RVΔG–UVI–E2-Crimson uptake yields true starter cells co-expressing EGFP and E2-Crimson (yellow), from which RVΔG spreads to presynaptic partners (red) carrying the same UVI. **(B)** The AAV starter construct encoding TVA^66T^::EGFP and N2cG under the CaMKII promoter. **(C)** The engineered RVΔG–UVI–E2-Crimson genome. A 14-bp pseudorandom UVI is inserted upstream of the M gene 3^*′*^ UTR. **(D)** Representative FACS plots of a pseudotyping control, Helper-GKo (abortive starters), and Helper (tracing) conditions. **(E)** FACS quantification of the proportion of E2-Crimson^+^ cells within viable singlets across conditions (Pseudotyping control, N=2, n=4; Helper-GKo, N=3, n=9; Helper/tracing N=4, n=13). **(F, G)** Proportion (F) and absolute number (G) of E2-Crimson^+^ cells per TVA^66T^ organoid (N=4, n=18), corresponding to healthy hESC-derived organoids in final tracing conditions. **(H)** Immunofluorescence images of traced organoids. Starter cells (AAV^+^/RVΔG– UVI^+^) and traced cells (RVΔG–UVI^+^) distribute across SATB2^+^, CTIP2^+^, and SCGN^+^ populations. Scale bars: 500 µm, 100 µm, 50 µm. **(I, J)** Rank-frequency distribution of UVIs supported by >2 reads, RVΔG–UVI–E2-Crimson library versus published datasets in full (I), and top 25 UVIs (J). **(K)** COSCO workflow schematic: AAV transduction, RVΔG–UVI infection, FACS isolation, scRNA-seq, and computational reconstruction. N=number of independent differentiation experiments. n=number of organoids. Box plots show median and IQR; whiskers indicate min to max.

Reliable network reconstruction required two additional optimizations: (1) a helper construct that maximized monosynaptic spread and restricted starter identity; and (2) a highly-complex, low-bias RVΔG-UVI-E2Crimson library to prevent spurious network merging (**Figure 1B–C)**. We first optimized the AAV helper to increase spread while restricting starter identity. All components of the helper construct were placed under the CaMKII promoter to preferentially trace from excitatory starter cells ^40,41^. The final helper construct, CaMKII-TVA^66T^::EGFP-p2a-N2cG (hereafter helper), packaged in the retrograde AAV (AAVrg) serotype, carries the TVA^66T^ variant to reduce abortive starter cell infections ^42^; simplifies the construct from tricistronic to bicistronic through a TVA^66T^::EGFP fusion; and shortens the 3^*′*^ regulatory region to reduce construct size from 4.9kb to 4.0kb **(Figure S1A–B**). These modifications substantially improved tracing outcomes (**Figure S1C–D**).

We next tested whether RV^+^ cells reflected true N2cG-dependent spread rather than nonspecific uptake. First, we showed that EnvA-pseudotyped rabies selectively enters cells expressing the TVA receptor (specificity = 99.985%; **Figure 1D)**. To quantify true tracing efficiency, we developed a dual-helper, FACS-based tracing assay that can selectively measure RV^+^ starter cells alone, or RV^+^ starter plus connected cells, in the same batch of organoids. We therefore generated an N2cG-knockout helper (helper-GKo), which permits starter-cell formation but abolishes transsynaptic spread **(Figure S1E)**. Matched helper/helper-GKo experiments confirmed that the majority of E2-Crimson^+^ cells are generated by N2cG-dependent spread (**Figure 1D–E; S1F**). We defined spread rate as the ratio of E2-Crimson^+^ cells in helper versus matched helper-GKo conditions and found that TVA^66T^ increased spread rate relative to TVA^WT^ from ~7:1 to ~17:1 (**Figure S1G**).

Sparse AAV labeling limits multi-starter/polysynaptic network merging by minimizing the chance that two or more starter cells are directly connected. Additionally, sparse RV labeling minimizes dual infection of starter cells. Observed labeling rates were 0.29% for AAV/GFP^+^ cells (putative starter cells) and 4.5% for RV/E2-Crimson^+^ cells (true starters and connected cells), translating to approximately 5,000–10,000 RV^+^ cells per organoid in final tracing conditions (**Figure 1F–G)**. Finally, dual-color rabies infection showed that multi-virion infection of individual cells was rare (<0.018% after Poisson modeling; **Figure S1H**, Methods). These observations show that dual AAV or RV infections were experimentally minimized, and any residual events were further removed computationally. Together, the final tracing design yielded improved tracing power with bright, sparse, excitatory-neuron-restricted starter populations **(Fig-ure 1H**).

We engineered a high-diversity RVΔG–UVI–E2-Crimson library using the CVS-N2c strain due to its improved tracing efficiency relative to other strains ^43^. A 14-bp pseudorandom, homopolymer-free UVI was inserted upstream of the M-gene’s 3^*′*^UTR, maximizing UVI recovery in 10x scRNA-seq ^44^ and yielding a theoretical complexity of 3.58 × 10^7^ UVIs (**Figure 1C**). Bulk sequencing of the purified viral library yielded 8.2 million reads, identifying 145,971 barcodes before and 25,707 after Shepherd sequence correction ^45^. The UVI library showed high barcode separation (median Hamming distance = 10; **Figure S1I–J**), minimizing error-driven misassignment, and markedly lower barcode representation bias than published libraries ^34,35,46^ (**Figure 1I–J; S1K–M**).

Fluorescence microscopy confirmed tracing across key cortical subtypes, including upper-layer excitatory (SATB2^+^), deep-layer excitatory (CTIP2^+^), and CGE-derived GABAergic interneurons (SCGN^+^) **(Figure 1H)**. Sparse EGFP^+^/E2-Crimson^+^ co-positive starter cells were distributed throughout the organoid and enriched in excitatory populations, supporting CaMKII promoter specificity (**Figure 1H**). Together, we established COSCO as a robust platform for large-scale connectomic mapping in human cerebral organoids (**Figure 1K**).

### COSCO reconstructs thousands of neuronal networks in cerebral organoids

To map the connectomic architecture of human cerebral organoids, we performed COSCO across 37 samples consisting of single- or small pools of 2–3 organoids, derived from six cell lines: two healthy stem cell lines (H9 hESC and 177 hiPSC), two TSC2^+/-^ hiPSC lines (TSC.p5.M and TSC.p24.M, hereafter “TSC”), and their isogenic healthy controls (TSC.p5.C and TSC.p24.C, hereafter: “Control”) expressing GCaMP6s, as described previously ^19^. All experiments used the same organoid differentiation protocol and viral system described above. Experiments were primarily conducted at 150 days *in vitro* (DIV), with a subset at 120 DIV to assess earlier developmental stages. Analyses included samples for all lines in Figure 2; all non-TSC (or wild type) lines in Figures 3; all lines in Figure 4; all 150 DIV non-TSC samples in Figure 5; and all TSC and isogenic control samples (all 150 DIV) in Figure 6.

**Fig. 2.**
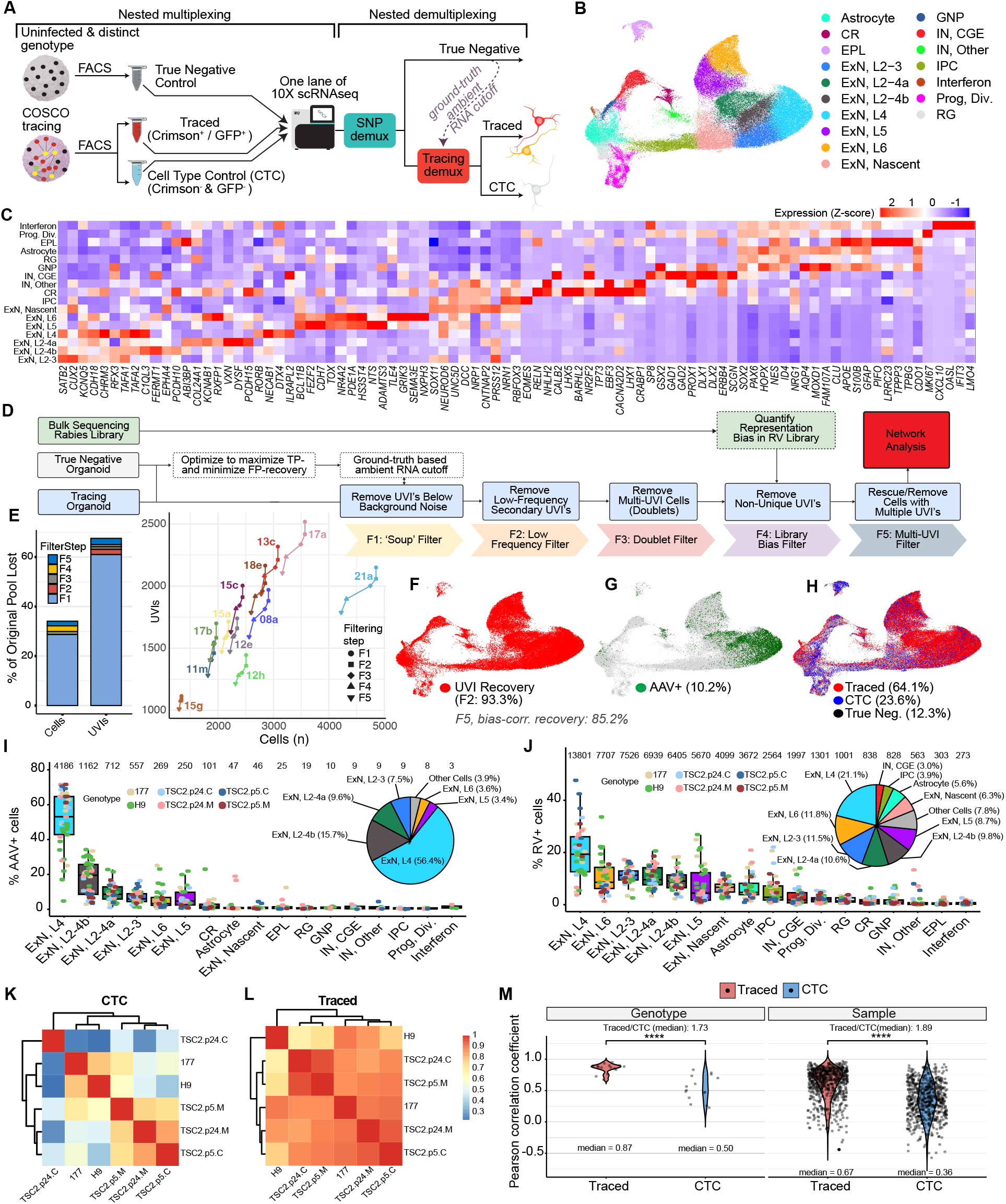
COSCO enables transcriptomic and connectivity profiling of single synaptic networks. **(A)** COSCO experimental paradigm. AAVrg- and RVΔG–UVI–E2-Crimson-transduced organoids were FACS-separated as: starter- and connected cells (EGFP^+^/E2-Crimson^+^ plus EGFP^−^/E2-Crimson^+^); CTC (EGFP^−^/E2-Crimson^−^; <10% of cells collected), and True Negative cells from a genetically distinct organoid, never exposed to any viruses. **(B)** UMAP of 113,757 high-quality cells annotated by cell state. **(C)** Heatmap of canonical marker genes confirming cell state assignments. Color indicates z-scaled expression level. **(D)** UVI identification and filtering pipeline overview. **(E)** Effect of sequential filtering steps (F1–F5) on UVI recovery across libraries. Left: relative loss from raw UVI^+^ cell counts highlight the importance of accurate ambient UVI correction (F1). Right: progression of a random sample of datasets through filtering steps F2–F5 in UVI-to-cell space shows dataset specific differences and various sources of errors. **(F)** UMAP highlighting cells with recovered UVIs (93.3% of Traced cells after F2 filtering). **(G)** UMAP highlighting AAV^+^ starter cells (10.2% of Traced, 0.94% of CTC cells and 0.25% of TN cells). **(H)** UMAP annotated by category post-filtering: Traced 64.1%, CTC 23.6%, True Negative 12.3%. **(I, J)** Distribution of starter (I) and connected (J) cells per sample across cell states, dots colored by genotype; numbers on top indicate total cells per cell state; pie charts show dataset-wide distribution. H9: N=5, n=13; 177: N=4, n=8; TSC2.p24.C: N=3, n=4; TSC2.p24.M: N=3, n=5; TSC2.p5.C: N=2, n=4; TSC2.p5.M: N=2, n=3. **(K, L)** Pearson correlation coefficients of CLR-transformed cell state proportions across genotypes in CTC and Traced populations. Heatmaps share a common color scale. **(M)** Distribution of correlation coefficients from (K-L and S2K-L). Each point represents one pairwise correlation value; median values and their ratios are labeled. Two-sided Wilcoxon rank-sum test. ^****^p < 0.0001. N = number of independent differentiation experiments. n = number of organoid samples.

**Fig. 3.**
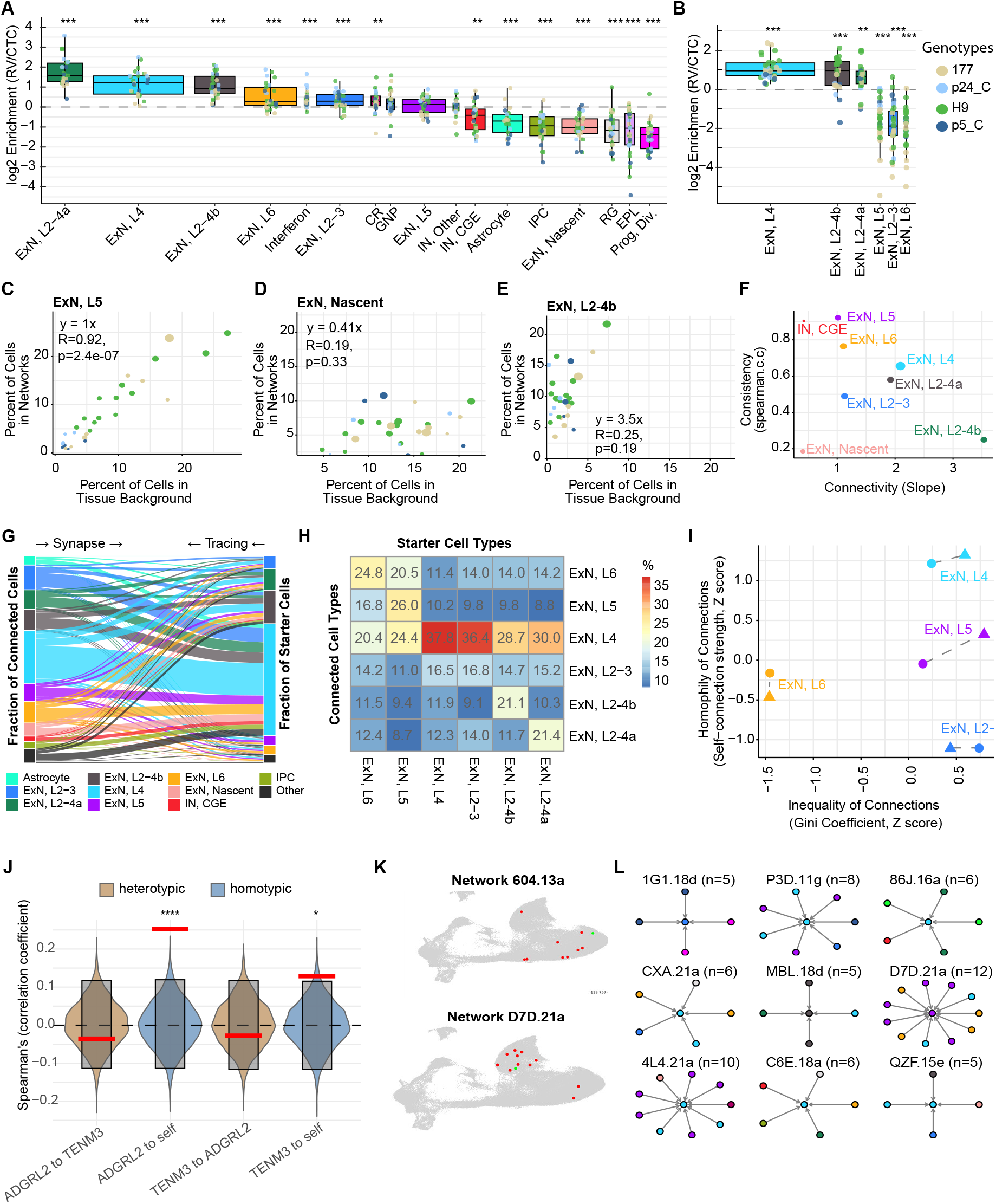
Cell-state-intrinsic wiring rules govern neuronal network assembly in human cerebral organoids. **(A)** Log2 fold Traced-to-CTC enrichment per cell state, across wild-type day 120–150 samples. Dots represent individual samples, colored by genotype; box width is proportional to CTC abundance; cell states ordered by mean enrichment BH adjusted MWW p-values: ^**^p < 0.01, ^***^p < 0.001, ^****^p < 0.0001. H9: N=5, n=13; 177: N=4, n=8; TSC2.p24.C: N=3, n=4; TSC2.p5.C: N=2, n=4. **(B)** Log2 fold enrichment of AAV^+^/RV^+^ (Trace) starter cells over AAV^−^/RV^−^ (CTC) cell states. Visual details, statistical testing and sample sizes as in (A). **(C–E)** Relationship between cell type abundance in Traced over CTC compartment across individual samples, colored by genotype and point size scaled by relative CTC abundance. Cell state connectivity (slope of the linear model) and consistency (Spearman correlation) quantified. **(F)** Connectivity and consistency values (as above) across neuronal subtypes. **(G)** Alluvial plot of relative cell state abundance and synaptic information flow across all UVI-defined networks pooled from all wild type organoids. Cell states below 2.5% abundance are grouped as “Other”. **(H)** Pairwise breakdown of input connectivity across neuronal subtypes. Percentages per starter cell type (columns) are shown. **(I)** Homotypic connectivity strength versus input inequality (Gini coefficient) in organoids (circles) and in mouse (triangles) across common excitatory neuronal subtypes. **(J)** Cross-cell correlation of *ADGRL2* and *TENM3* expression in two-cell networks (one starter, one connected cell; red) against a 10,000-permutation null (violins; grey boxes: 95% CI). Spearman correlations computed for homotypic and heterotypic pairs; empirical significance assessed against the null (of shuffled starter identities). ^*^p < 0.05, ^****^p < 0.0001. **(K, L)** Representative reconstructed networks on UMAP (K); and a random sample of networks (5–12 cells) as graphs, colored by cell type; (L). N = number of independent differentiation experiments. n = number of scRNA-seq samples.

**Fig. 4.**
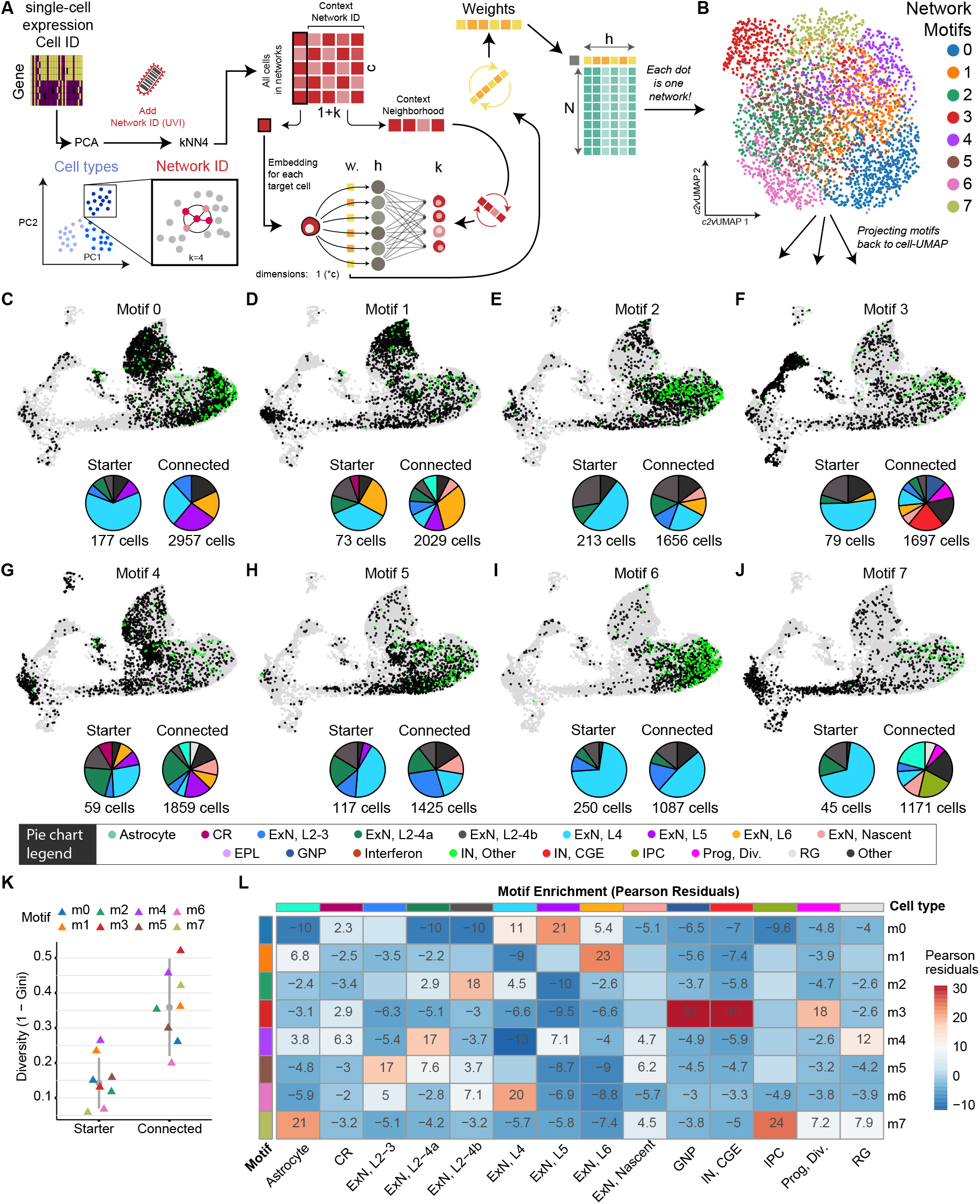
Recurrent synaptic network motifs revealed by clone2vec embedding. **(A)** clone2vec converts kNN neighborhoods (in transcriptional PCA space) into network co-occurrence relationships, then embed all networks so that ones with similar network-contexts appear proximal. For clarity, the schematic shows k=4; the analysis used k=15. **(B)** Network-UMAP calculated from the clone2vec (c2v) embedding, colored Leiden clustering of networks; each point represents one UVI-defined network. **(C–J)** UMAPs of all cells per network motif (m0–m7). Starter cells in green, connected cells in black, all other cells in gray. Pie charts show cell type composition of starter-(left) and connected cells (right) per motif; total cell counts below. Cell states below 5% abundance are grouped into “Other” (black). **(K)** Cell type diversity (1 − Gini coefficient) of starter and connected cells across network motifs (triangles). Grey vertical lines indicate the IQR, grey dot the median. **(L)** Heatmap of Pearson residuals across network motifs in wild-type DIV120–DIV150 single-starter networks. Red and blue indicate over- and under-representation compared to a chi-square independence model; values beyond *±*2 are shown. Tiles with both observed and expected counts below 10 were masked.

**Fig. 5.**
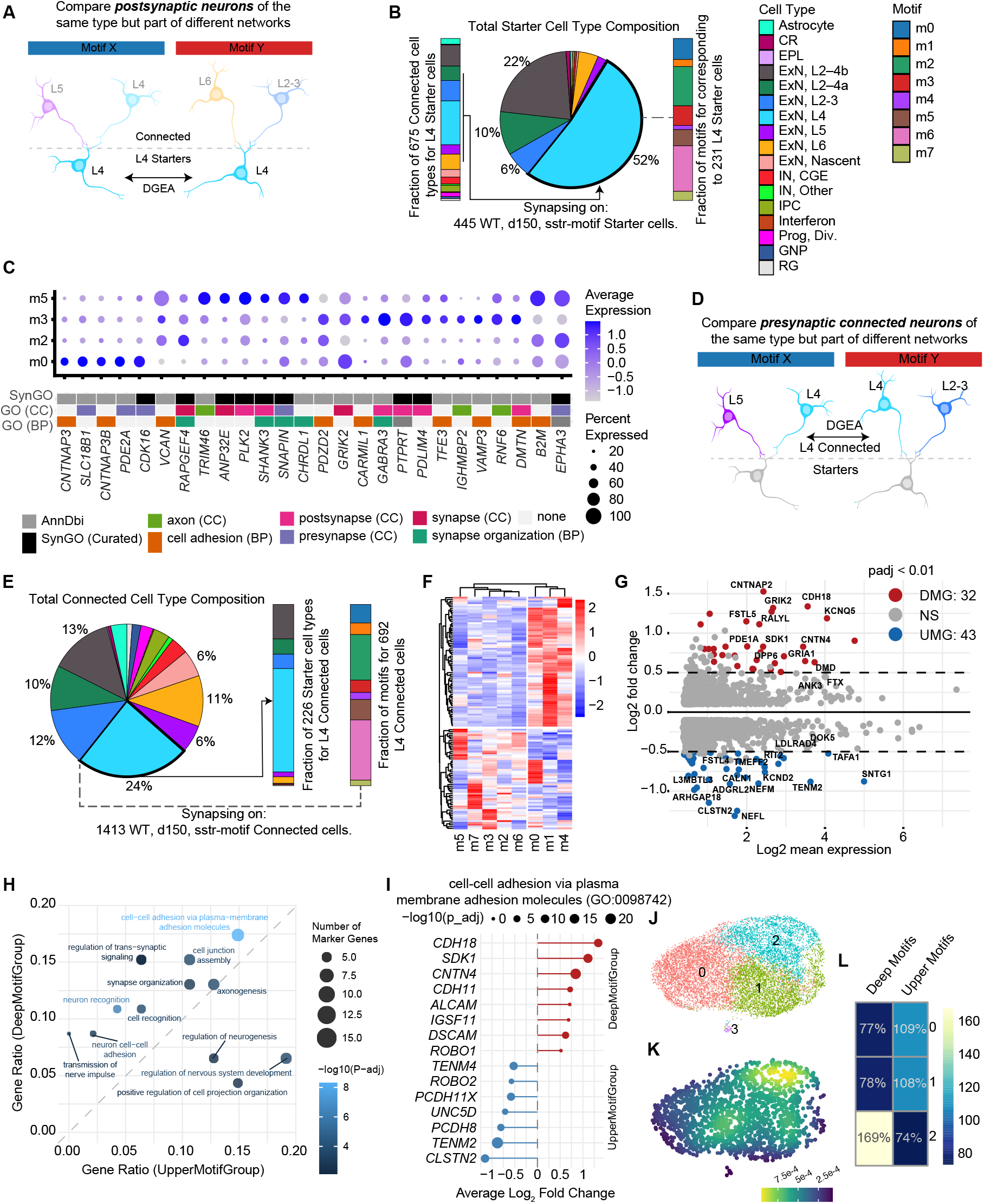
Synaptic network context is reflected in neuronal gene expression programs. **(A)** Conceptual framework for DGEA within postsynaptic L4 starter neurons across network motifs. **(B)** Distribution of motif-assigned starter cells from wild-type DIV150 single-starter UVI-defined networks (n = 445). Pie chart: cell-state composition of starter cells, with L4 highlighted (n = 231; 52%). Bar plot 1: cell-state distribution of connected cells synapsing onto L4 starter cells (n = 675). Bar plot 2: motif distribution of the same L4 starter cells used for DGEA. **(C)** Synapse relevant motif-specific marker genes of L4 starter cells from m0, m2, m3 and m5, annotated GO and SynGO synaptic terms (BH-adjusted MWW test; p < 0.01, min.pct = 0.5, log2FC > 1). **(D)** Conceptual framework for DGEA within L4 connected cells across network motifs. **(E)** Distribution of motif-assigned L4 connected cells from wild-type DIV150 UVI-defined networks with zero or one detected starter cell (n = 1,413). Pie chart: cell-state composition of the connected-cell subset, L4 connected cells highlighted. Bar plot 1: cell-state distribution of detected starter cells within the above single-starter networks. Bar plot 2: motif distribution of the same L4 connected cells used for DGEA. **(F)** Hierarchical clustering of marker genes in L4 connected cells from motif-wise one-versus-all DGEA (n = 1,413; BH-adjusted MWW test; p < 0.01, min.pct = 0.5, log2FC > 0.5), revealing a Deep Motif Group (m0, m1, m4) and an Upper Motif Group (m2, m3, m5, m6, m7). **(G)** MA plot of pairwise differential expression between Deep and Upper Motif Groups in L4 connected cells (n = 1,413 cells; 32 DMG-associated and 43 UMG-associated genes; absolute log2FC > 0.5, adjusted p < 0.01). **(H)** GO term enrichment comparison between Deep and Upper Motif Groups. Each point represents a GO category; x- and y-axis positions indicate the fraction of marker genes in those terms found in Deep or Upper Motif Groups; size indicates gene count; color indicates global GO enrichment significance (−log10 BH-adjusted p-value). **(I)** Top differentially expressed genes in GO:0098742 ranked by average log2FC. Point size indicates significance (−log10 BH adjusted DGEA p-value); direction indicates Deep versus Upper Motif Group association. **(J)** Transcriptomic subclustering of wild-type L4 ExNs. **(K)** Weighted kernel density projection of Deep Motif Group membership onto the transcriptomic L4 UMAP. **(L)** Enrichment of Deep and Upper Motif Group neurons across three transcriptomic subclusters 0–2, relative to their overall abundance among motif-assigned L4 neurons (100% = expected).

**Fig. 6.**
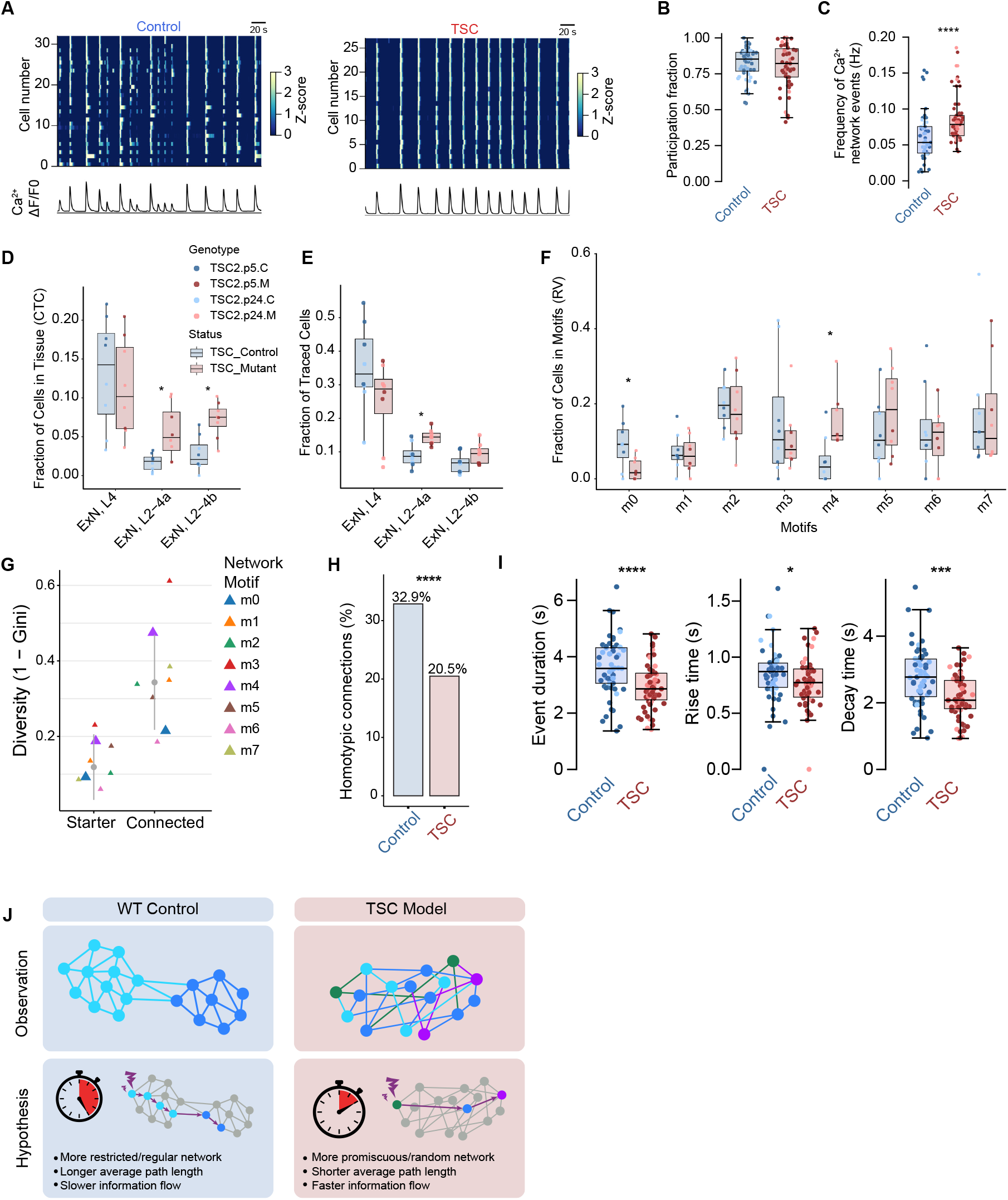
TSC drives aberrant, heterogeneous connectivity, linked to upper layer cell type switch and network hyperactivity. **(A)** Representative GCaMP6s recordings from Control and TSC organoids. Raster plots show calcium transients per neuron (rows); traces (bottom) show the aggregate fluorescence across the field of view. **(B)** Fraction of detected active neurons participating in synchronous network events in Control and TSC organoids. Each point represents one field of view (FOV). Control: TSC2.p24.C: N=6 batches, n=11 organoids, 21 FOV; TSC2.p5.C: N=7, n=15, 36 FOV. TSC: TSC2.p24.M: N=5, n=6, 13 FOV; TSC2.p5.M: N=7, n=18, 44 FOV. Whiskers extend to 1.5*×* IQR. Two-sided Wilcoxon rank-sum test. **(C)** Frequency of spontaneous synchronous network events. Sample sizes as in (B). Two-sided Wilcoxon rank-sum test. ^****^p < 0.0001. **(D, E)** Distribution of selected upper-layer excitatory neuron states in Control versus TSC organoids. Cell populations: CTC controls (D) and Traced cells (E). Fractions were calculated per sample, denoted by points, colored by genotype. TSC2.p24.C: N = 3, n = 4; TSC2.p24.M: N = 3, n = 5; TSC2.p5.C: N = 2, n = 4; TSC2.p5.M: N = 2, n = 3. FDR-adjusted ^*^p < 0.05, two-sided median permutation test on per-sample fractions. **(F)** Distribution of traced cells across network motifs in Control and TSC organoids. Fractions were calculated per sample across motifs m0–m7, denoted by points, colored by genotype. Sample sizes and statistics as in (D-E). **(G)** Cell state diversity of starter and connected TSC cells across network motifs (triangles), as in Figure 4K. Grey vertical lines indicate the IQR, grey dot the median. M0 and M4 are highlighted, see (F). **(H)** Average fraction of homotypic connections (same cell state) among starter-connected cell pairs in Control and TSC organoids. Two-sided Fisher’s exact test. ^****^p < 0.0001. **(I)** Calcium transient kinetics: event duration, rise time, and decay time. Sample sizes as in (B). Two-sided Wilcoxon rank-sum test. ^*^p < 0.05, ^***^p<0.001, ^****^p < 0.0001. **(J)** Model of TSC-associated network reorganization. Top row: observed increase in network heterogeneity in TSC. Bottom row: hypothesized link between altered topology and altered functional activity. N=number of independent differentiation experiments. scRNA-seq: n=number of demultiplexed samples. Calcium imaging: n=number of organoids; FOV=number of fields of view recorded.

To minimize technical bias and reduce library costs by ~40%, we developed “Nested Multiplexing”, an experimental-computational strategy **(Figure 2A)**. For each experiment, we pooled three samples into a single 10x library: (1) the Crimson^+^ “Traced”-fraction comprising helper-and-tracer^+^ starter cells and tracer^+^ connected cells; (2) the Crimson^-^ cell-type composition control (“CTC”), sorted from the same organoids; and (3) the uninfected cells from a genomically distinct organoid never exposed to viral constructs (“True Negative” or “TN”). Following WGS/WES-based SNP demultiplexing, True Negative cells served as a library-specific ground-truth estimate of ambient viral RNA, enabling precise separation of Traced from CTC cells, where the CTC population defined the sample-specific tissue composition. In addition, we leveraged TNs for two other purposes: separate AAV^+^ from AAV^-^ cells, and a parameter optimization for UVI selection (Methods).

Across 37 experiments, each comprising Traced, CTC, and TN populations, cells were processed through standard 10x library construction, Illumina sequencing, and 10x Cell Ranger preprocessing, recovering 219,650 cells in total. We applied stringent quality controls using a custom pipeline built on Seurat v5^47^, removing low-quality cells, stressed cells using Gruffi ^48^, and mispatterned cells, identified by projection onto an organoid reference atlas ^49^. We demultiplexed True Negative, Traced and CTC samples with a novel open-source R package, NestedMultiplexer (Methods), removed genotypic multiplets ^50^, and integrated datasets using Harmony ^51^. After subsequent QC filtering, we retained 113,757 high-quality single cells.

To account for transitional populations in developing organoids, we refer to transcriptionally defined populations as cell states throughout. Combining canonical markers with label transfer from fetal and organoid datasets ^51–53^, we identified 17 major cell states across 86,817 neurons, 16,651 progenitors, and 10,289 glial cells **(Figure 2B–C; S2A–B)**. Non-neuronal populations included dividing progenitors (Prog, Div.; *KI67*+), radial glia (RG; *SOX2*^+^), intermediate progenitor cells (IPC; *EOMES*^+^), glial neuronal progenitors (GNP; *GSX2*^+^/*DLX1*^+^/*S100B*^*+*^/*AQP4*^+^), astrocytes (*AQP4*^+^/*GFAP*^+^/*S100B*^*+*^), and ependymal-like cells (EPL; *S100B*^*+*^/*PIFO*^+^/*TPPP3*^+^), as well as the smallest, progenitor-like cluster (298 cells), specifically expressing antiviral, NF-kappaB- and TNF-pathway related genes (Interferon; *IFIT3*^*+*^*/CXCL10*^*+*^*/OASL*^*+*^). Reported CVS-N2c-induced genes ^54^ (*RIGI[DDX58]*^*+*^*/IFIT1*^*+*^*/MX1*^*+*^*/IL6*^*+*^) were largely absent from neurons and concentrated in this cluster. We additionally detected CGE-derived interneurons (IN, CGE; *NR2F2*^+^/*SCGN*^+^/*GAD1*^*+*^) and Cajal-Retzius cells (CR; *RELN*^+^/*LHX5*^+^). Excitatory lineages included nascent excitatory neurons (ExN, Nascent; *NRP1*^+^/*SOX11*^+^) and cells spanning canonical cortical layer identities: ExN, L2–3 (*SATB2*^+^/*CUX2*^+^), ExN, L4 (*RORB*^+^/*NECAB1*^+^), ExN, L5 (*BCL11B*^+^/*FEZF2*^+^), and ExN, L6 (*FOXP2*^+^/*TLE4*^+^) ^10,55–58^. We additionally identified two upper-layer excitatory cell states (ExN, L2–4a and ExN, L2–4b) by shared marker expression with L2–4 ExNs and validated this by label transfer from fetal atlases **(Figure S2C; Table S1)** 52,59–61. These populations were distinguished by high expression of distinct protocadherins (*PCDH15* and *PCDH10*, respectively), which are implicated in synaptic connectivity and neurodevelopmental disorders ^62^ **(Figure S2B)**. Cell-state distributions were broadly consistent across genotypes in both CTC and Traced fractions, with expected enrichment of mature neuronal states in the latter and comparable sample contributions within each genotype, indicating that downstream analyses are not confounded by compositional or batch effects **(Figure S2D–G)**.

We next asked whether rabies infection itself introduces a confounding transcriptional signature, as reported *in vivo* ^63^. We found a limited impact validated by two methods: the separation in UMAP and PCA spaces; and cell-state specific changes in gene expression. RV^+^ (Traced) and RV^-^ (CTC) cells remained intermixed across transcriptomic embeddings, with mean cell-state-specific median LISI scores ^51^ of 1.86 (theoretical range, 1–2) in Harmony-UMAP (CV, 4%), 1.68 in Harmony Components (CV, 6%), and 1.56 in non-integrated PCA space (CV, 7%; **Table S2**). Given its limited effect, it was not necessary to regress out infection status during integration, arguing against a dominant rabies-induced transcriptional state. Next, we measured rabies-associated root-mean-square (RMS) log2 fold changes in gene expression between Traced and CTC cells. Differences were markedly smaller than differences between cell states and variation between cell lines within the same cell state **(Figure S2H)**. Thus, rabies tracing largely preserves transcriptomic identity and is unlikely to affect connectivity-associated gene expression. We speculate that this muted effect may be due to the lack of microglia and systemic immune response in organoids.

To reconstruct high-fidelity synaptic networks, we developed a computational pipeline that systematically corrects for ambient viral RNA, library bias, and other technical artifacts **(Figure 2D–E)**. The primary challenge in barcoded single-cell tracing experiments is the elimination of ambient viral barcode RNA, which can lead to false network assignments. For UVI calling, we again leveraged the nested multiplexing design: True Negative cells were used to perform a parameter optimization step to identify the optimal read- and Unique Molecular Identifier (umi)-depth thresholds for UVIs in every library, maximizing true-positive- and minimizing false-positive detection (F0), after which UVIs falling below those library-specific thresholds were removed (F1). Low-frequency UVIs (<20% of top UVI-umis) in cells with multiple barcodes were removed, counteracting the residual false positive signal from the percentile-based F1 filtering. These rare barcodes were scarcely detected in connected partners (F2). Transcriptomic multiplets were then removed to prevent spurious network merging (F3), and library bias was corrected by comparing observed UVIs against the expected number of times a UVI would occur, based on its frequency in the viral library and on the estimate of the number of starter cells in that sample (F4). The probability of identical UVIs independently infecting multiple neurons was then simulated, and any barcode with a >5% probability of recurring was blacklisted; remaining multi-UVI conflicts were conservatively resolved to finalize network assignments (F5). We define a network as all cells sharing a UVI after F5 filtering. Across the full dataset, this pipeline recovered a UVI in 93.3% of Traced cells at F2 and finally retained 85.2% (n=62,126) after library bias correction at F5 **(Figure 2F; S2I–J)**. We identified 10.2% (n=7,422) of traced cells as starters, by detecting AAV transcripts at FPR=0.0025 (**Figure 2G**). Traced, CTC, and True Negative cells comprised 64.1%, 23.6%, and 12.3% (Traced=72,909, CTC=26,854, TN=13,994) of total cells in the dataset, respectively **(Figure 2H)**. The CaMKII promoter-controlled helper efficiently targeted excitatory neurons, with starter cells comprising 96.8% excitatory neurons, including 90.1% upper-layer and 56.4% L4 excitatory neurons, while connected cells were broadly distributed across mature excitatory and inhibitory states **(Figure 2I–J)**, consistent with diverse presynaptic inputs.

A salient feature of the dataset was the reproducibility of synaptic network organization across organoids. As described elsewhere, cellular composition of organoid tissue (CTC) can vary substantially across cell lines and samples ^64^ (**Figure 2K; S2K**). In contrast, Traced-cell composition was highly correlated across cell lines and samples (**Figure 2K– M; S2L**), indicating that variability in tissue composition was not propagated into network composition. Thus, synaptic networks of excitatory starter cells assemble into reproducible architectures, despite differences in the available cell-state landscape. This demonstrates that organoid network organization is intrinsic to the assembling neurons rather than imposed by the surrounding tissue.

### Cell-intrinsic wiring rules govern neuronal networks in cerebral organoids

Having validated COSCO across barcode recovery, starter-cell detection, transcriptomic fidelity, and reproducibility, we turned to the biological organization of networks. We first examined the cell-state specificity of tracing, and found that mature neuronal populations, particularly upper-layer excitatory neurons, were enriched among Traced cells relative to CTC cells (**Figure 3A**). As expected, given the lack of a functional postsynaptic machinery, nascent excitatory neurons; late-born, immature interneurons; glial and progenitor cells were depleted. Because glial and progenitor cells were present in networks, albeit depleted, we examined this phenomenon further, finding that astrocytes and IPCs accounted for the majority of the RV^+^ glia and progenitor annotations (**Figures S3A–B**). Astrocytes co-localize with neurons and often participate in tripartite synapses, and were previously traced by rabies ^46^ and therefore this residual immature and glial signal is consistent with other studies ^34,65,66^. We next analyzed cell state frequencies in starter cells (min. 100 cells per cell state) in the same way and found that most upper-layer neurons were >1.8-fold enriched among true starter cells (**Figure 3B**). Taken together, these results demonstrate that COSCO tracing predominantly captures synaptically mature neuronal cell states.

To determine whether network participation simply mirrors tissue abundance, we compared, per sample, each cell state’s frequency in the Traced fraction to its matching CTC fraction (tissue background). We then quantified connectivity (linear-regression slope) and consistency (Spearman correlation) in a linear model across cell states **(Figure 3C–E; S3C– G)**. We found that connectivity patterns were strongly cell-state dependent (**Figure 3F**). Deep-layer excitatory neurons (L5, L6) exhibited lower connectivity and high consistency, whereas nascent excitatory neurons and CGE-interneurons showed weak connectivity, as expected given their immaturity ^67,68^. By contrast, L2–4b excitatory neurons displayed high connectivity but low consistency, indicating that their incorporation into networks is regulated by additional factors, and tissue abundance is a poor predictor of their network abundance. We then moved from population-level connectivity to barcoded network analysis. We recovered 10,510 unique networks, of which 34% contained more than 2 cells, and 82% were distributed across the four wild-type lines. Starter cells were recovered in 17% of networks, and 2.6% contained more than one starter cell; these were excluded from downstream analyses (**Figure S3H–J**). We then investigated network organization at the level of pre- and postsynaptic pairs. Published neuron-to-neuron connectivity-and-transcriptome mapping studies have recovered presynaptic networks for at most hundreds of single postsynaptic neurons ^34–36^. Measured on the same terms, COSCO recovered networks for 1,779 starter cells (8–59x fold increase) and 3,263 connected cells (4–10x fold increase) **(Figure 3G)**.

Having established the scale of network recovery, we next asked how connectivity is organized between starter and connected cells. We could analyze this relationship on the population level due to the large number of networks with starter cells recovered. We calculated the fractional innervation of major excitatory neuron types (**Figure 3H**), which revealed two organizing principles. First, most neuronal cell states preferentially connect to the same cell types (homotypic connections, or homophily). Second, deep-layer and upper-layer excitatory neurons exhibited distinct input architectures: deep-layer starter cells received diverse inputs from multiple cell states, whereas upper-layer starter cells exhibited more selective input patterns, dominated by L4 excitatory neurons. Besides homophily, we quantified input diversity as 1-Gini coefficient, a measure of compositional evenness that captures diversity when measured within a fixed set of categories (cell states). Together, homotypic connectivity and input diversity resolved three excitatory input signatures, most clearly represented by L4, L2/3 and L6 ExNs (**Figure S3K–L**). To assess whether these wiring patterns reflect intrinsic neuronal properties rather than organoid-specific effects, we compared the above connectivity to layer-specific RVΔG tracing data from adult mouse visual cortex ^69^. Importantly, apart from the biological model and the absence of viral barcoding, this dataset employed matching experimental conditions, namely RV strain, scRNA-seq workflow, and annotation strategy (**Figure S3M–N**). Remarkably, relative connectivity patterns were conserved across systems. Both *in vivo* and *in vitro*, L6 ExNs exhibited the highest input diversity and L4 ExNs showed the strongest homophily with low input diversity. L2/3 ExNs displayed low homophily connectivity with similarly restricted input diversity (**Figure 3I**). Thus, the layer-specific connectivity signatures observed *in vitro* mirror those of the mammalian cortex *in vivo*.

Cell-surface recognition molecules play a key role in neuronal partner selection. We therefore asked whether known receptor-ligand matching predicts partner choice. We focused on *ADGRL2* (Latrophilin-2) and *TENM3* (Teneurin-3), two major candidates organizing perinatal mouse circuit assembly through homophilic molecular interactions and correlated topographic expression maps ^70,71^. Across 2,242 connected or starter-connected pairs, *ADGRL2*-*ADGRL2* and *TENM3*-*TENM3* showed significant expression cross-correlation, whereas *ADGRL2*-*TENM3* cross-correlations were insignificant (**Figure 3J**). Our correlative analysis cannot explain context-dependent molecular processes ^72^, nor the complex spatial, temporal and cell-type specific expression patterns ^73^. Nonetheless, it shows that human synaptic networks *in vitro* recapitulate developmental mouse wiring patterns, such as the Tenm3 *→* Tenm3 and Adgrl2 *→*Adgrl2 rule ^71^. We next moved from pre-aggregated analysis to unbiased, label-free single-network analysis which revealed diverse, structured connectivity patterns, spanning selective neuronal cell states **(Figure 3K–L)**. This raised the question of whether these patterns robustly occur across the full populations of networks.

### Unbiased network embedding uncovers recurrent patterns of connectivity

After analyzing connectivity by predefined cell states, we next applied clone2vec to identify recurrent network patterns without predefined cell-state annotations ^38^. clone2vec is an adaptation of word2vec ^74^ and embeds groups of cells (networks) based on the transcriptomic neighborhoods of their constituent cells ^38^. Although originally developed for barcoded lineage tracing, clone2vec operates on predefined groups of cells, and is thus also applicable to traced networks. Briefly, clone2vec calculates barcode co-occurrences within k-nearest-neighborhoods of each cell in networks, which are then embedded into a low-dimensional (c2v) space using a skip-gram model that factorizes the co-occurrence structure by optimizing an ePCA objective with stochastic gradient descent (**Figure 4A; S4A**). This produces an embedding of networks, which is followed by Leiden clustering and _**c2v**_UMAP visualization.

We applied clone2vec to 2,969 networks containing ≥ 3 cells, which resolved eight distinct network clusters (hereafter “motifs”; **Figure 4B**). These motifs incorporate networks both with and without a recovered starter cell. As 96.8% of starter cells were excitatory neurons, network origin can be confidently assigned to them even when starter cells are not present. Projection of each motif’s members onto the cellular UMAP revealed distinct compositions of starter- and connected-cell states for each motif (**Figure 4C– J; S4B–C**). To assess whether motif structure reflects biological organization rather than stochastic barcode spread, we generated randomized control datasets by shuffling UVIs across cells and recomputed network embeddings. Experimental networks were significantly dissimilar from all randomized controls (P = 5^*^10^-8^ to 1.3^*^10^-4^), which did not significantly differ from one another (**Figure S4D**). Thus, motif structure is not explained by random labeling or local transcriptomic similarity alone. Consistent with a biological basis for the embedding, 120 and 150 DIV networks separated along the X-axis of the _**c2v**_UMAP, revealing an age-associated axis of network organization, while samples otherwise appeared well mixed (**Figure S4E–F**). These early-stage sam-ples, which constituted a minority of our data, were included to probe whether network organization is sensitive to developmental stage, rather than to perform a systematic analysis of network maturation. Nonetheless, we reasoned that strong age-associated separation might capture the sequential maturation of excitatory neuron classes. Since deep-layer neurons mature before upper-layer neurons, younger organoids should contain networks with greater deep-layer participation ^57^. Indeed, deep-layer neurons (L5/L6) were ~20% more abundant in 120 DIV networks than in 150 DIV networks (**Figure S4G**), and 120 DIV networks preferentially occupied deep-layer-enriched motifs (m0, m1, and m4; **Figure 4C–D, 4G**). Thus, even this limited dataset recapitulates the developmental shift from early deep-layer enriched networks toward later upper-layer dominated networks, consistent with the later emergence of upper-layer neurons over developmental time.

Cellular diversity varied across motifs and differed between starter and connected cells, which revealed two features of motif organization **(Figure 4K)**. First, connected-cell populations were consistently more diverse than starter-cell populations, with median diversity values of ~0.35 for connected cells and ~0.15 for starter cells. Second, starter-cell diversity did not determine connected cell diversity. For example, while diversity in m6 was restricted in both populations, m3 combined low starter diversity with high connected-cell diversity. This restriction in m6 reflects the selective dominance of L4 ExNs in both starter and connected cells. This suggests that the previously observed aggregate L4 ExN homophily (**Figure 3H–I**), arises from a distinct subset of L4 ExNs, rather than broadly characterizing all L4 ExN-starter networks.

We then calculated cell-state enrichments across all motifs, using Pearson residuals to quantify deviations from expected cell-state frequencies. We found that within individual motifs, typically a single cell state is dominant, with an additional 2–5 cell states enriched (rows), whereas each cell state is enriched across 1–3 motifs (columns; **Figure 4L**). When cell state enrichment per motif was stratified by starter cell state, we found that connected cell composition correlated with motif identity and not with starter cell identity **(Figure S4H)**. Individual starter cell states also contributed to multiple motifs. Starter cells annotated as the same cell state (e.g., L4 ExN) therefore participated in networks of different composition. This indicated that current transcriptomically defined cell-state annotations do not capture the features that specify network connectivity. Among possible explanations, scRNA-seq analysis pipelines may overlook transcriptional differences that predict connectivity. We used COSCO to test whether such differences exist.

### Network architecture is reflected in gene expression

A central prediction of COSCO is that neuronal connectivity should be reflected in transcriptional programs at the single-cell level. We therefore asked whether neurons belonging to different motifs exhibit corresponding differences in gene expression. To identify gene expression patterns that correlate with connectivity rather than cell-state or age differences, we restricted our analyses to L4 ExNs in wild-type, 150 DIV organoids.

We first examined how gene expression in L4 ExN starters correlates with their presynaptic connectivity, as defined by motif identity (**Figure 5A**). After selecting L4 ExN starter cells across network motifs, we retained motifs with >20 L4 ExN starters and performed differential gene expression analysis (DGEA) between them (**Figures 5B; S5A– B**). The resulting markers included canonical genes of cell adhesion, neurotransmission, as well as novel candidates for synapse formation and organization. To better interpret these patterns, we focused on genes with known synapse- and cell adhesion-related functions using Gene Ontology (GO) and SynGO ^75^ terms (**Figure 5C**). Because this analysis was deliberately restricted to L4 ExN starters, these motif-associated signatures should be interpreted as fine-scale connectivity-linked differences within one cell state, rather than the extensive differences defining cell states. Here, motif 3 provided a clear example of how presynaptic network context is reflected in L4 ExN starter-cell gene expression. This motif was selectively enriched for presynaptic *SCGN*^+^ CGE interneurons, and it contained 79% of those innervating any L4 ExN starters (**Figure S5C–D**). Correspondingly, L4 ExN starters in m3 preferentially expressed *GABRA3*, encoding the α3 subunit of the GABA_A_ receptor. This match between presynaptic inhibitory input and postsynaptic *GABRA3* expression provides a direct example of connectivity-associated gene expression, while corroborating the biological validity of the clone2vec-defined motifs. Beyond this expected neurotransmitter–receptor match, GABA_A_ receptors directly bind presynaptic neurexins ^76^, and concordantly we found that *NRXN3* was selectively expressed in the presynaptic CGE interneurons.

Motif 5 (m5) provided another example of connectivity-linked gene expression. m5 L4 ExN starter cells received inputs enriched for nascent and recently born L2–3 ExNs, the latest-born excitatory populations in our organoids **(Figure 4H)**. As developing neurons use synaptic scaling to stabilize activity while incorporating new excitatory connections ^77,78^, we examined whether m5 starter cells expressed the corresponding machinery **(Figure 5C)**. Indeed, they specifically upregulated the established synaptic-scaling regulators *PLK2* ^79^ and *SHANK3* ^80^. Thus, m5 couples developmentally young excitatory inputs to the postsynaptic transcriptional programs expected to stabilize these connections. Together, these results show that motif assignment and transcriptional profiles correspond to expected synaptic relationships. While most genes identified here have established cellular (synaptic or adhesion-related) functions, our results suggest that their role may be also understood by their impact on the network level, which so far largely remained uncharacterized.

We next asked whether connected neurons (as opposed to starter neurons) exhibit similar connectivity-defined transcriptional differences, again focusing on L4 ExNs (**Figure 5D–E**). Using the same DGEA framework, we identified motif-associated markers that reflect the composition of their synaptic networks. These motif-associated markers segregated into two higher-order groups, defined by clustering marker gene coexpression (m0, m1, m4 versus m2, m3, m5, m6, m7; **Figure 5F**). The first group collectively accounted for 71% of all connected deep-layer (L5/L6) neurons and was therefore termed Deep Motif Group (DMG). The second group comprised motifs generally lacking deep-layer neurons and was therefore termed Upper Motif Group (UMG) (**Figure 4C–J)**. By collapsing motifs into these two groups, we could test whether L4 ExN gene expression differs according to deep-layer co-wiring with greater statistical power then looking at individual motifs (**Figures 5G and S5E**).

We next calculated GO Biological Process (GO-BP) enrichments for either motif group. L4 ExNs in the DMG were preferentially enriched in adhesion- and synaptic-transmission-related terms, while L4 ExNs in the UMG were more enriched in development and maturation-related terms (**Figures S5F–H**), potentially reflecting developmental differences, as upper-layer cortical circuits emerge later during corticogenesis both *in vivo* and in organoid models ^58,81^. Because several terms were enriched in both groups, we next examined the underlying gene usage. For shared enriched terms, gene-ratio plots revealed a more nuanced picture of this trend (**Figure 5H**). For example, even when the same plasma-membrane adhesion term was enriched in both motif groups, the underlying genes differed. DMG enrichment was driven mainly by cadherin/contactin/IgSF-related recognition genes (*CDH18, CDH11*/*CNTN4*/*SDK1, ALCAM, IGSF11, DSCAM, ROBO1*), consistent with a cell-surface recognition program dominated by molecules capable of homophilic and identity-matching interactions. In contrast, UMG enrichment involved teneurin/protocadherin/calsyntenin and guidance-receptor genes (*TENM2, TENM4*/*PCDH8, PCDH11X*/*CLSTN2*/*ROBO2, UNC5D*). Thus, the same ontology label captured distinct molecular strategies: one centered on cell-identity recognition, the other on adhesion, guidance, and synaptic refinement **(Figure 5I)**. Similar divergence was observed across other shared GO terms **(Figure S5I)**.

Finally, we asked whether motif-defined connectivity states are also present in purely transcriptomic space but remain uninterpretable without connectomic context. To address this, we re-clustered all 150 DIV wild-type L4 ExNs, regardless of tracing status, using a standard, unsupervised Seurat workflow. This expanded dataset increased the total number of L4 ExNs and, correspondingly, statistical power. We identified three main subclusters, 0–2 **(Figure 5J)**. Standard single-cell analysis interprets clusters using DGEA followed by GO enrichment. For transcriptionally similar sub-clusters, however, this approach returns largely the same terms, leaving the subclusters indistinguishable, as was also the case here. We therefore projected connectivity data onto this UMAP and found that deep-layer connectivity is highly enriched in subcluster 2 (**Figure 5K–L**). By linking transcriptional separation to deep layer connectivity, we provide a tangible interpretation for the cluster identity. As examples, we show marker genes with known or putative synap-tic roles, co-segregating with deep-layer connectivity **(Figure S5J)**. COSCO therefore enables the (re)interpretation of standard single-cell results, and allows researchers to exploit the full, statistically robust, transcriptome-wide marker set to decipher the synaptic code. Together, these results demonstrate that neuronal connectivity, at both the starter and connected cell level, is coordinated with transcriptional programs at single-cell resolution. Combining connectomic and transcriptomic data, we recapitulated known principles of synaptic organization and revealed a previously uncharacterized, connectivity-linked molecular diversity in L4 ExNs.

### Increased heterogeneity, reduced homophily and a cell-state shift in TSC

After characterizing network formation in healthy controls, we asked whether COSCO could identify connectivity defects in TSC, a neurodevelopmental disorder caused by heterozygous loss-of-function variants in either *TSC1* or *TSC2*. These mutations lead to hyperactivated mTOR signaling which, in the brain, results in developmental cortical lesions, cortical tubers, autism spectrum disorder phenotypes, and early-onset drug-resistant epilepsy in up to 90% of patients ^21^. Previous work in our group using TSC organoids recapitulated key morphological and electrophysiological disease features, including dysmorphic neurons, tumorigenesis, network hyperexcitability, and epileptiform high-frequency oscillations ^19,20^. However, how TSC alters the excitatory network architecture at the single-neuron level remains unknown. To understand the changes in TSC, we analyzed organoids from two patient-derived hiPSC lines (*TSC2*^+/-^; TSC.p5.M and TSC.p24.M, “TSC”), along with their isogenic controls (*TSC2*^+/+^; TSC.p5.C and TSC.p24.C, “Control”). All lines express a genetically encoded calcium indicator (GCaMP6s) to verify the functional phenotype in DIV 150 organoids. Both Control and TSC organoids exhibited reproducible, spontaneous synchronous calcium activity (network events) and comparably high event participation rates **(Figure 6A–B, S6A)**. However, TSC organoids showed significantly higher event frequency, consistent with network hyperexcitability observed at later time points ^19^ **(Figure 6C)**.

To test whether structural changes in connectivity could underlie this functional phenotype, we performed COSCO on 16 organoids derived from 10 independent batches, yielding 3,223 CTC and 17,404 traced TSC cells; and 5,609 CTC and 16,020 traced Control cells, organized into 1,840 and 1,323 networks, respectively **(Figure S6B)**. We first investigated if TSC organoids show changes in tissue composition (CTC) and found that they exhibited a significant increase in L2– 4a and L2–4b ExNs and a decrease in L4 ExNs **(Figure 6D; S6C)**. Traced cells in TSC organoids showed a similar increase in L2–4a and L2–4b ExNs, and a decrease in L4 ExNs **(Figure 6E;S6D)**. Importantly, the combined upper-layer excitatory pool of L2–4a, L2–4b, and L4 neurons in networks did not change in TSC (**Figure S6E**; L2-3 showed no significant difference). Thus, in organoids, TSC disrupts the balance of upper-layer excitatory cell states participating in networks.

To determine the consequences of cell-state redistribution for network organization, we analyzed Control and TSC cells across clone2vec network motifs. TSC organoids showed a significant network redistribution from motif 0 to motif 4, which displayed the highest proportion of L2–4a cells among all motifs **(Figure 6F)**, consistent with, but not fully explained by the above-described cell-state redistribution. In fact, it also reflected broader differences in network organization. The TSC-depleted motif 0 represented a cell-state-restricted and stereotyped architecture, whereas motif 4 networks spanned more neuronal cell states, and showed less constrained (or more promiscuous) connectivity across those **(Figure 6G)**. To validate this pattern independently of clone2vec analysis, we quantified homotypic connectivity across all reconstructed pre- and postsynaptic cell pairs. This showed a 38% relative decrease in within-cell-type-connectivity in TSC organoids, corroborating that synaptic partners in TSC are more heterogeneous in cellular composition **(Figure 6H)**. Finally, we analyzed individual networks (n≥ 5) traced from L4 ExNs or from any starter cell and again found a trend toward increased heterogeneity in TSC **(Figure S6F–G)**. Thus, TSC shifts networks toward less constrained motif architectures, revealing disease-associated rewiring beyond changes in cell-state composition.

Classical network science models show that reducing connectivity constraints can shorten average path length and predict faster information propagation dynamics ^82–84^. While these models describe loss of constraint in physical space, we observed a conceptually analogous loss of constraint in cell-state space, with TSC neurons receiving fewer homotypic inputs and participating in more heterogeneous networks. Although these models are too simplistic to fully capture neural dynamics, for networks composed predominantly of excitatory neurons, they provide a useful framework for inferring the direction of changes in network-dynamics parameters. In short, topological changes toward less restricted wiring in TSC would be expected to favor faster population-level Ca^2+^ dynamics. In line with this expectation, TSC organoids showed shorter rise time, decay time, and duration of synchronous Ca^2+^ events **(Figure 6I)**. While this interpretation remains a conceptual framework, it supports the idea that network topology can contribute to the early hyperactive phenotype in TSC **(Figure 6J)**, and that COSCO can resolve functionally relevant disease-associated changes in neuronal network architecture.

Together, these results establish COSCO as a scalable approach to resolve the synaptic and transcriptional architecture of human neuronal networks. In cerebral organoids, COSCO reveals reproducible, cell-state-associated principles of network self-organization and shows that these principles are disrupted in TSC, aligning with observed Ca^2+^ dynamics.

## Discussion

Here, we have described COSCO, a method to map synaptic networks in human cerebral organoids at single-cell transcriptomic resolution. By combining barcoded rabies-based viral tracing, scRNA-seq, nested multiplexing, optimized UVI processing, and network embedding, COSCO reconstructs more than 10,000 networks while preserving cell-state and transcriptional information. This scale allows network organization to be studied as a quantitative phenotype across large populations, rather than in isolated instances. We analyzed connectivity at multiple complementary levels, each suited to a specific biological question and optimized for statistical power so that all fractions of the data contributed to conclusions, rather than only the smaller subset of networks containing starter cells. Starter-connected pair analysis resolved directional relationships and allowed comparisons to mouse data. Co-embedding networks with and without starter cells revealed recurrent motifs, while starter-defined networks supported detailed gene-expression-connectivity analyses.

Cerebral organoids are reductionist models that have been shown to recapitulate key aspects of human fetal brain development which make them a tractable system to study principles of human neuronal network formation ^8,17,85–87^. Here, we demonstrate that cerebral organoids form biologically relevant and reproducible neuronal networks. We show that organoid networks are not random nor simply reflections of tissue composition but are shaped by consistent wiring preferences. Networks maintained a stable and defined cell type composition despite tissue variability across samples, suggesting robust principles of network assembly. We further show that excitatory input signatures recapitulate patterns observed in the mouse visual cortex ^69^, and that connectivity-associated gene expression is consistent with *in vivo* observations ^71^. Thus, cerebral organoids preserve fundamental aspects of cortical circuit organization, providing a foundation for studies of human network assembly, under conditions not accessible *in vivo*.

We next demonstrate that self-organization is a key principle of neuronal network formation in cerebral organoids. This is supported by several observations. First, cerebral organoids develop under stable, insulated, and spatially unconstrained conditions, without input from sensory systems or other brain regions, favoring intrinsic self-organization over external instruction. Second, the selectivity and robustness of network assembly in absence of external guidance suggest a self-organizing mechanism. Third, synaptic partner selection followed cell-state-specific rules, exemplified by distinct patterns of homophily and input diversity. Fourth, clone2vec revealed recurrent network motifs, showing how simple pairwise connectivity preferences between cell states can give rise to higher-order network organization. Fifth, known molecular regulators of wiring *in vivo* show the expected connectivity-to-expression relationships in organoids, as discussed below.

A second principle emerging from our organoid data is that core elements of connectivity are encoded in molecular state. This link was most evident in L4 ExNs, where the same nominally homogeneous cell type separated according to network context. Postsynaptic starter neurons assigned to different motifs expressed distinct transcriptional programs, linking presynaptic input architecture to postsynaptic gene expression. This was illustrated by the association of post-synaptic *GABRA3* expression with CGE interneuron inputs, and by homotypic *ADGRL2* and *TENM3* expression across connected pairs, extending anatomical-scale observations in the mouse brain to human microcircuits ^70,71^.

The principle was further supported in connected L4 ExNs, which formed two motif-groups distinguished both by deep-layer connectivity and distinct adhesion and guidance programs. When projected onto an unsupervised reclustering of L4 ExNs, DMG localized to a single subcluster, linking otherwise unexplained transcriptomic differences to connectivity. Because this approach co-clustered all L4 ExNs in the whole dataset, it extended the analysis beyond directly traced cells and yielded a large, robust set of candidate genes. These results do not establish causality, but they demonstrate that COSCO can reveal molecular differences that standard scRNA-seq analysis alone cannot resolve, highlighting candidate connectivity-associated genes and molecular programs for future study.

Finally, in TSC patient-derived organoids, we show that organizational principles of network formation are disrupted in disease, and that these changes may help explain early electrophysiological phenotypes, before late-connecting interneurons’ dysfunction becomes dominant ^19,20^. Our data suggest that TSC alters the composition of the upper-layer excitatory neuron pool and shifts their wiring toward less constrained and more heterogeneous patterns. Reduced wiring constraints in theoretical network models can lead to increased global connectivity and accelerated signal propagation ^82,84,88^, providing one possible framework for interpreting hyperexcitability. However, TSC is a multifactorial disorder, and altered network topology is likely only one of several contributing mechanisms. Within these limits, COSCO links *TSC2* mutation to changes in cell state, network topology, and activity.

COSCO complements existing connectomic approaches by trading spatial and subcellular resolution for scale and transcriptomic depth. Electron and expansion microscopy preserve synaptic-resolution structure in individual samples, whereas COSCO resolves network composition at scale. It also links connectivity to transcriptional programs, identifies recurrent connection patterns, enables studying of the largely inaccessible human fetal network development, and how network organization changes in disease. COSCO also differs from viral projectome mapping and spatial methods, which primarily target anatomical mapping *in vivo*, but face trade-offs in gene coverage, throughput, resolution, or cost ^31,35,37,89,90^. It is conceptually closest to the pivotal SBARRO approach ^34^, and a recent synaptosome- and nuclei-seq-based approach using molecular tracers, which increases transcriptomic fidelity at the cost of experimental complexity ^36^. Altogether, COSCO achieves the scale and analytical precision that enables fundamental insights into human neuronal network organization.

Future applications should expand starter-cell targeting, extend COSCO to other disease models, larger cohorts, regionalized or guided organoid protocols, incorporate addi-tional interneuron and glial classes, map developmental trajectories across organoid ages, or extend to chimeroids and assembloids. Integrating connectivity with spatial transcriptomics, calcium imaging, and electrophysiology, particularly pairing activity and connectivity measurements from matched organoids, should be especially powerful for linking network structure to function.

### Limitations of the study

This study has several limitations. Cerebral organoids allow intrinsic human network self-organization to be studied in a controlled system, but they model only a subset of the processes that shape connectivity *in vivo*. They lack sensory-driven and long-range circuit interactions, vascularization, and full anatomical organization. In addition, batch, line, and maturation variability cannot be fully controlled or regressed out.

Also, COSCO measures a selected fraction of connectivity. It selectively maps monosynaptic retrograde inputs onto virally defined starter cells, not multi-step circuits, synapse number or strength. Our CaMKII-driven AAVrg helper design selects for upper-layer excitatory starter cells, prioritizing interpretability over comprehensiveness. This allowed us to interpret networks without starter cells, but limited access to other populations like inhibitory-starter circuits. Our conclusions are also drawn from a single organoid differentiation protocol and predominantly one developmental time point (150 DIV). Whether conclusions translate to other protocols and developmental stages remains untested. Additionally, while we have shown that rabies infection had little transcriptional impact in immune-deficient organoids, we still restricted our transcriptional analysis to comparing RV^+^ populations. Even so, we cannot fully exclude that a subset of connectivity-associated transcripts reflects aspects of viral infection.

Sequencing readout trades anatomy for scale. Spatial position, morphology, and subcellular structures are lost. Cell loss during single-cell preparation reduces apparent network size in a stochastic, experiment-dependent manner. Consequently, absolute network sizes cannot be measured, nor reliably compared across samples. Cell loss further leads to numerous orphan networks missing a starter cell, which in previous approaches lacking latent embedding, could not be analyzed. Advances in low pressure microfluidic isolation and gentler library preparation methods already mitigate this problem.

Finally, COSCO links connectivity, gene expression, and disease-associated network changes, but it does not establish causality. It identifies transcriptional programs correlated with synaptic partner selection but does not prove that these genes instruct wiring. In TSC organoids, we describe the consequences of *TSC2*^*+/-*^ mutation: hyperexcitability, cell-state redistribution, and motif shifts, but cannot determine causality between these phenotypes.

## Resource availability

### Lead contact

Requests for further information and resources should be directed to and will be fulfilled by the lead contact, Juergen A. Knoblich

### Materials and Data availability

All plasmids will be available from the authors upon request. Whole genome sequencing (WGS) data used for cell-line genotype assignment are available under SRX3471959/SRR6377128 for hESC H9; EGAD00001006333 for hiPSC TSC.p5.M; EGA50000002008 for TSC.p24.M; EGAD00001008769 for hiPSC lines 176.1 and 177.8, with access managed through and [IMBA iPSC Biobank Webshop](https://shop.vbc.ac.at/ipsc_biobank). Raw sequencing data for single-cell experiments is available under: EGA50000002010. Processed single cell data is available for reviewers and will be publicly available upon acceptance.

### Code availability

The code for barcode-, single-cell- and electrophysiology analyses, as well as tools developed for analyzing single-cell connectomics data is accessible to reviewers, and COSCO analysis tools will be publicly available on GitHub, upon acceptance. The following custom function libraries were used for the analysis: Stringendo, ReadWriter, CodeAndRoll2, MarkdownHelpers, ggExpress, Seurat.Utils, all freely available under github.com/vertesy

## Acknowledgements

Special thanks to Yoav Ben-Simon (Allen Institute) and Flavia Leite at the ISTA viral core for initial discussions, plasmids and viruses; Nina Corsini, Peter Karchenko, Amit Zeisel and Zhige Lin (Technion) for technical discussions and help; Noelia Urbán and Katherina Taverini (IMBA) for discussion on glial lineages; Julius Brennecke and Christoph Bock for advising throughout the process; Mark Noble, Noelia Urbán, Oliver Eichmüller, Emilia Asgrimsdottir, Sakurako Wong, Nina Corsini and Julian Scherer for careful revision of the manuscript (IMBA); among others, Gerald Schmauss, Mario Nezhyba and Marietta Weninger at BioOptics- and Thomas Grentzinger, Philip Wolff and Alex Vogt at the NGS Core Facilities of the Vienna Bio-center (VBC) for their outstanding work and commitment. For iPSC generation and sequencing, we thank the IMBA iPSC Biobank and the VBCF NGS facilities. The computational results presented were obtained using the CLIP cluster (https://clip.science).

## Author contributions

Conceptualization, Á.V., R.N., B.D., and J.A.K.; methodology, Á.V., R.N., B.D., C.V.L.; investigation, Á.V., R.N., B.D; stem cell and organoid culture, R.N., B.D., C.Z., S.K; formal analysis, Á.V., R.N., M.N., T.R.B, M.Z., S.I; software, Á.V., S.I, M.Z., M.N., T.R.B; resources, J.A.K.; data curation, R.N., Á.V.; visualization, Á.V., R.N., M.Z.; writing – original draft, Á.V., R.N.; writing – review & editing, all au-thors; supervision, J.A.K., I.A.; funding acquisition, J.A.K., I.A., Á.V., R.N., M.Z.

## Declarations

### Declaration of generative AI and AI-assisted technologies in the writing process

During the preparation of this work the author(s) used various LLMs for rewriting, formatting and editing text and for writing, editing, refactoring, and debugging code. After using these tools and services, the authors reviewed and edited the content as needed and take full responsibility for the content of the published article.

## Methods

### Experimental Method Details

#### hESC and iPSC generation

The healthy control hESC line (H9; WA09, female) was acquired from WiCell. The healthy control hiPSC lines (176; 176J clone#1, female and 177; 177J clone#8, female) were generated at the IMBA Stem Cell Core Facility and are part of the IPSC Biobank. After informed consent was obtained, a skin biopsy was taken and fibroblasts were isolated for iPSC reprogramming using the Sendai virus (CytoTune-iPS 2.0 Sendai Reprogramming Kit, Thermo Fisher Scientific) carrying the Yamanaka reprogramming factors OCT3/4, SOX2, c-MYC, and KLF4. TSC patient iPSC lines and their isogenic controls were originally described elsewhere ^20^, and a GCaMP6s construct under the human synapsin promoter was integrated into the AAVS1 locus as described previously ^19^ (TSC2.p5.C= TSCp5-30s-hSyn1-GCaMP6s cl.2; TSC2.p5.M= TSCp5-32s-hSyn1-GCaMP6s cl.3; TSC2.p24.C= TSCp24#4 R.2C4 hSyn1-GCaMP6s cl.4; TSC2.p24.M= TSCp24#4 hSyn1-GCaMP6s cl.9). The use of all cell lines was approved by the local ethics committee of the Medical University of Vienna (MUV), and informed consent was obtained from all participants. All experiments involving human cell lines, particularly the ones involving human embryonic stem cells, were conducted in agreement with Austrian law and regulations.

#### Maintenance and passaging of hESCs and iPSCs

All cells were cultured on 6-well plates (Corning, Cat no. 3516) coated with hESC-qualified Matrigel (Corning, Cat. no. 354277) and maintained in complete mTeSR™1 medium (STEMCELL Technologies, Cat no. 85875). Cells were fed daily with 2mL of mTeSR™1 (feeding with 4mL allowed skipping of one feeding day per week) and passaged after reaching 60-80% confluency (every 3–5 days). For passaging, cells were exposed to 0.5mM EDTA (Thermo Fisher Scientific, Cat no. 15575020) diluted in DPBS^-/-^ (Gibco, Cat no. 14190-250) for 3 minutes at 37°C. After this incubation period, EDTA was removed and 1mL mTeSR™1 was added to the cells by gently spraying the bottom of the well. The cells were then triturated into small clusters. Cells in solution were then transferred to newly coated Matrigel 6-well plates at ratios of 1:8 to 1:15 to continue expanding. Cells were maintained at 5% CO_2_ at 37°C. All cell lines were authenticated using a short tandem repeat (STR) assay, tested for genomic integrity using single-nucleotide polymorphism array genotyping and routinely tested for mycoplasma ^91^. All cell lines were used within 10 passages of the last STR profiling.

#### Organoid generation

Human iPSCs and hESCs were grown to 60–80% confluency. On day 0, cells were dissociated into a single-cell suspension by Accutase (Sigma-Aldrich, Cat no. A6964-100ML) treatment for 5 minutes at 37°C, followed by manual trituration. Cells were seeded in an ultra-low binding 96-well plate (Corning, Cat no. COR7007) at a density of 9,000 live cells/well, in 150µL of complete mTeSR™1 medium with 10µM ROCK inhibitor (Y-27632 2HCl) (Selleck Chemicals, Cat no. S1049). To facilitate embryoid body (EB) formation cells were centrifuged in the 96-well plate for 2 minutes at 120g. On day 2 90% of media was aspirated and replaced with complete mTeSR™1 medium. This was repeated on day 4. On day 5, 90% of media was aspirated and replaced with Neural Induction (NI) medium prepared as follows (recipe for 500mL): DMEM/F12 (1:1) 1x, with Hepes: 500mL (Thermo Fisher Scientific, Cat no. 11330-057); N2 supplement: 5mL (Thermo Fisher Scientific, Cat no. 17502048); Glutamax supplement-I (100X): 5mL (Thermo Fisher Scientific, Cat no. 35050-038); MEM Non-essential amino acid (NEAA) solution (100x): 5mL (Thermo Fisher Scientific, Cat no. 11140050); Antibiotic-Antimycotic (100x): 5mL (Thermo Fisher Scientific, Cat no. 15240062). On days 7 and 9, 90% of media was aspirated and replaced with NI. On day 10, EBs were removed from the 96-well plate with a 200µL Wide Bore Filtered Pipette Tip (Thermo Fisher Scientific, Cat no. 2069GPK) and placed onto a parafilm strip (Merck, Cat no. HS234526B) with indentations manually formed by pressing into the top an empty pipette box. Excess medium was manually removed. EBs were then embedded in 20µL Matrigel droplets (Corning, Cat no. 354234) and allowed to solidify by incubating at 37°C for 25–30 min. After this incubation period, embedded organoids were gently removed from the parafilm strip and placed into 10cm dishes (TPP, Cat no. Z707651) in 20mL Differentiation Medium without vitamin A (Diff-A), prepared as follows (recipe for 500 mL): DMEM/F12(1:1) 1x, with Hepes: 250 mL; Neurobasal: 250 mL (Thermo Fisher Scientific, Cat no. 21103049); N2 supplement: 5 mL; B27 minus Vitamin A supplement: 10 mL (Thermo Fisher Scientific, Cat no. 12587010); Recombinant human Insulin in solution: 125 µL (Merck, Cat no. I9278); Glutamax supplement: 5 mL; MEM Non-essential amino acid solution (100x): 2.5 mL; Antibiotic-Antimycotic (100x): 5 mL. On Day 13, 90% of Diff-A medium was removed and replaced with Differentiation Medium with vitamin A (Diff+A), prepared as follows (recipe for 500 mL): DMEM/F12(1:1) 1x, with Hepes: 250 mL; Neurobasal: 250 mL; N2 supplement: 5 mL; B27 plus Vitamin A supplement: 10 mL (Thermo Fisher Scientific, Cat no. 17504044); Recombinant human Insulin in solution: 125 µL; Glutamax supplement: 5 mL; MEM Non-essential amino acid solution (100x): 2.5 mL; Antibiotic-Antimycotic (100x): 5 mL; Sodium bicarbonate: 500mg (Sigma-Aldrich, Cat no. S5761). The organoids were then placed on an orbital shaker (InforsHT, Celltron) within the 37°C incubator rotating at 57 RPM. From day 16 to day 40, every 3 days, 90% of media was aspirated from organoids and replaced with Diff+A medium. On Day 40, organoids were fed with 75% Diff+A and 25% complete BrainPhys neuronal medium. 500 mL complete BrainPhys neuronal medium was prepared as follows (recipe for 500 mL): BrainPhys basal medium: 475 mL (STEMCELL Technologies, Cat no. 05790); B27 plus Vitamin A supplement: 10 mL (Thermo Fisher Scientific, Cat no. 17504044); N2 supplement: 5 mL (Thermo Fisher Scientific, Cat no. 17502048); Ascorbic Acid: 2.5 µL (200 nM, Sigma-Aldrich, Cat no. A4544-25G); CD Lipid Concentrate: 1 mL (Thermo Fisher Scientific, Cat no. 11905031); Antibiotic-Antimycotic (100x): 5 mL (Thermo Fisher Scientific, Cat no. 15240062); 20% Glucose solution: 3.4 mL (10 mM, made in house). This was supplemented with brain-derived neurotrophic factor (BDNF) (20 ng/mL; STEMCELL Technologies, Cat no. 78005.3), glial cell line-derived neurotrophic factor (GDNF) (20 ng/mL; STEMCELL Technologies, Cat no. 78058.3), and bucladesine sodium (db-cAMP) (1 mM; MedChemExpress, Cat no. HY-B0764), added fresh before feeding. On day 43 this was repeated with 50% Diff+A and 50% complete BrainPhys medium with supplements. On day 46 this was repeated with 25% Diff+A and 75% complete BrainPhys medium with supplements. From day 50– 150, feeding was performed with 100% complete BrainPhys medium with supplements. On days 60, 90, 120 and 150, organoid medium was supplemented with 1% Matrigel diluted into the medium to supplement the growth and stability of the organoids.

#### AAV cloning

All cloning was performed with NEBuilder® HiFi DNA Assembly Master Mix (NEB, Cat no. E2621L) as per the manufacturer’s instructions. Briefly, overhangs of 20–60 bp were designed for insertion, and vectors were ei-ther amplified using KOD One™ PCR Master Mix (Merck, Cat no. KMM-101NV) or digested using the appropriate restriction enzymes. The plasmids resulting from cloning were purified using SPRIselect beads (Beckman Coulter, Cat no. B23318) and transformed into Stbl3 chemically competent bacteria (Thermo Fisher Scientific, Cat no. C737303). The bacteria were plated on agar plates with appropriate selection media. Colonies were picked and amplified in LB media with appropriate antibiotics, followed by miniprep (NEB, Cat no. T2010S) and submission for whole plasmid sequencing by Plasmidsaurus. The AAV TVA^WT^ helper construct was used as described previously (Addgene #219638) ^92^. To introduce 66T and GKo mutations, primers with the relevant point mutations were used to amplify TVA and G respectively, followed by Gibson assembly as described above. Combinations of AAV cargos containing TVA^66T^ and G were prepared (4 combinations in total: TVA^WT^/G (Addgene #219638), TVA^66T^/G (Addgene #259763), TVA^WT^/GKo (Addgene #259761) and TVA^66T^/GKo (Addgene #259762)) to be able to measure and compare spread for both TVAs.

#### AAV generation

Adeno-Associated Virus was generated based on previous protocols ^93,94^. Briefly, HEK293T cells were grown to 80% confluency in 10×15cm^2^ dishes. On Day 0, cells were then transfected overnight with 20mL DMEM+10%FBS (Cytiva, Cat no. SH30071.03HI) with plasmids carrying the capsid for AAV-Retro (gift from ISTA), helper genes (Addgene #112867) and cargo genes (described above) according to the calculations described in the literature ^94^. For the transfection reagent, linear, 25,000MW Polyethylenimine was used (Polysciences, cat. no. 23966-1). On Day 1, the media on all plates was replaced with 20mL Advanced DMEM (Fisher Scientific, Cat no. 12-491-023) + 5% FBS. On Day 4, all media was harvested, stored at 4°C and replaced with fresh 20mL Advanced DMEM.

The purification of AAV was based on previous work with minor modifications ^93^. On Day 5, Triton-X100 (Sigma Aldrich, 1003339793) was added to the viral media harvested on Day 4, up to 1%. 20mL of the harvested media was then added back to the 15cm^2^ plates. The plates were placed on a shaker (50rpm) and were incubated at 37°C for 1 hour to detach and lyse cells. The media from each plate was then harvested and centrifuged for 30 minutes at 6000g in an Avanti J-26 XP (Beckman Coulter) centrifuge to remove larger cellular debris and aggregates. The media was then filtered using a 0.22µm PES filter (Sartorius, Cat no. VF20P7). Next, the virus-containing, clarified media was concentrated by tangential flow filtration using a peristaltic pump (Master-Flex Easy-load., Cat no. 07514-10) and a 100,000MW cutoff PES filter (Sartorius Cat no. VF20P4). The media was concentrated down to 5mL and the filtration system was flushed with 25mL of PBS.

To purify AAV virions from the concentrated media, an iodixanol gradient was used along with PBS-MK (26.3 mg of MgCl_2_ and 14.91 mg of KCl in 1 × PBS in a final volume of 100 mL) or PBS-MK + NaCl (100mL of PBS-MK + 5.94g of NaCl). 15mL of viral media was layered on the bottom of a 25×89mm polypropylene tube (Beckman Coulter, 326823) using a syringe and a long, blunt tip needle. Layers of OptiPrep (StemCell, Cat no. 00222) were then injected underneath the media, slowly raising it to the top and filling the tube. The volumes and composition of the layers were in order: 8mL of 15% Iodixanol (2.25mL of 60% Iodixanol stock (STEMCELL Technologies, Cat no. 00222) + 6.7 mL of 1M PBS-MK with Sodium Chloride (Sigma-Aldrich, Cat no. S9888)), 6mL of 25% iodixanol (3mL of Iodixanol stock + 4.2mL of PBS-MK), 5mL of 40% Iodixanol (4mL of Iodixanol stock + 2mL of PBS-MK) and 5mL of 60% Iodixanol stock. To distinguish between gradients, 40µL of phenol red (Sigma-Aldrich, Cat no. P0290) was added to the 25% and 60% layers. The tubes were sealed and placed in a Beckman Coulter 70ti ultracentrifuge rotor and were centrifuged for 2 hours at 44.400rpm at 18°C. The tubes were then removed from the rotor, pierced using an 18-gauge needle from the bottom and the 40% layer was collected by gravitational flow. To wash out the Iodixanol and further concentrate the virus, the collected layer was diluted to 50mL using PBS, filtered using Steriflip (EMD Millipore, Cat no. SE1M003M00) and then concentrated using Vivaspin 20 100,000MW PES filters (Sartorius, Cat no. VS2041) down to approximately 300µL (100µL per 15cm^2^ dish). The viral preparation was aliquoted and stored at −70°C. qPCR was used to titer the virus as described previously ^95^.

#### Rabies plasmid library barcoding

To barcode the CVS-RVΔG genome at the 3’ end of the M Gene, first, two PaqCI restriction sites compatible with Golden Gate ligation were cloned into the viral genome using the following insert primer: 5’-gtctgaagaggacaaagactcttctctgcttctagaataactaggaggctgGCAGGTGagnctnctga CACCTGCtcagattatgcgcctgcaagtgtatcacttgtttacctctggaggagagagca-3’. To incorporate unique barcodes, oligos with the following PaqCI compatible landing sites were designed: 5’-CACGTGAAGACCAGCTAGC CCACCTGCcagcggagHNBNVNDNHNBNVNattactgaGC AGGTGCGCCCGTATCTGTCTTCGTC-3’. To amplify the barcode-oligos, qPCR was performed using AmpliTAQ 360 (Thermo Fisher Scientific, Cat no. 4398886) and EvaGreen (Biotium, Cat no. 31000). Specifically, 0.2µL of 5µM Bar-coded Oligo Template was amplified using 10µL each of 5µM of forward and reverse primers in 50µL reaction volume with 2.5µL of EvaGreen added. Reactions were set up to cycle as per manufacturer’s instructions. The reaction was stopped when the log-phase began (around Cycle 7), and the newly synthesized, double-stranded barcode fragments were purified using Ampure XP (Beckman Coulter, Cat no. A63881) following manufacturer’s instructions.

After the synthesis of double-stranded DNA from the single-stranded oligos above, Golden Gate ligation was performed with a pre-digestion step. First, 250ng of RV plasmid was digested using 1µL of PaqCI (NEB, Cat no. R0745L) and 1µL of PaqCI Activator in rCutSmart Buffer for 4 hours at 37°C, followed by inactivation for 20 minutes at 72°C. The cut vector was then purified using SPRIselect Beads (Beckman Coulter, Cat no. B23318) at 0.6x concentration. Golden Gate assembly was then performed in 50µL reaction volume using 100ng of cut vector, 1:2 ratio of barcode-oligo, 1µL of T4 DNA ligase (NEB, M0202M), 1µL of PaqCI, 1µL of PaqCI activator, 2µL of T4 ligase buffer and Milli-Q H_2_O. The reaction was cycled using the following program: Step 1: 37°C for 5 mins, Step 2. 22°C for 5 mins Step 3: repeat from Step 1. 12x. Step 4. 37°C for 30 mins, Step 5. 75°C for 20 mins. The now barcoded RV genome library was then purified from the reaction tube using SPRIselect Beads at 0.6x concentration and used for amplification.

To amplify the library, MegaX DH10B™ T1R Electro-comp™ Cells (Thermo Fisher Scientific, C640003) were electroporated using GenePulser Xcell (BioRad) according to the manufacturer’s instructions. Cells were plated on 15cm^2^ agar plates with Ampicillin and were left to incubate overnight. The following day, cells were washed off using ice-cold LB media and were shaken for 1 hour at 37°C. Cells were then spun down at 6000g for 15 mins, weighed and purified using the Qiagen GigaPrep kit (Qiagen, 12123). This plasmid library was then used to produce the barcoded rabies viral libraries.

#### Rabies virus generation

Pseudotyped rabies virus was produced as previously described ^96^. To produce the barcoded rabies virus library, existing protocols ^34^ were adopted with modifications. The virus-producing HEK293T-G-T7 (Gift from ISTA), described in ^96^, was generated by transducing HEK293T cells (Sigma-Aldrich, Cat no. 12022001) with Lentivirus made from MMLV-CAG-SADB19_oG-IRES-Puro (Addgene, Cat no. 172367) and MMLV-CAG-G_oT7pol-IRES-BSD (Addgene, Cat no. 172369). Cells were grown and maintained in culture media (DMEM+10% FBS) with Puromycin (3µg/mL) (Sigma, Cat no. P9620-10ML) and Blasticidin (15µg/mL) (Sigma, Cat no. SBR00022-10ML) added.

To begin production, two T225 plates (Thermo Fisher, Cat no. 159933) of HEK293T-G-T7 were grown until 60-70% confluency. On Day 0, the media on cells was refreshed with 45mL of DMEM+10%FBS and the plates were transfected at a 4:1:1:1:1 ratio by the RabV Library plasmid (131.36µg/T225 flask), pCAG-B19N (Addgene, Cat no. 59924, 32.84µg/T225 flask), pCAG-B19P (Addgene, Cat no. 59925, 32.84µg/T225 flask), pCAG-B19L (Addgene, Cat no. 59922, 32.84µg/T225 flask) and pCAG-T7pol (Addgene, Cat no. 59926, 32.84µg/T225 flask). For transfection, Takara Xfect (Takara Bio, Cat no. 631318) was used as per manufacturer’s instructions. Cells were transfected for 6h at 37°C, after which media was replaced with Production Media (Advanced DMEM, with 5% FBS, 1x Anti-Anti and 1x GlutaMAX added) and cells were moved to 35°C. Between Day 1 and Day 6, media was replaced daily with 45mL of Production Media. Visible clusters of RV+ cells were apparent from Day 2. On Day 7, media was not replaced, but rather 45mL of extra Production Media was added. On Day 8, the ~90mL of media from each plate was harvested, pooled, filtered using a 0.22µm filter and was pseudotyped.

To pseudotype the RV library produced in the previous steps, BHK-EnvA cells (Gift from IST, ^97^) were grown to 80-90% confluency in Production Media + Puromycin in 15cm^2^ dishes. 90mL of filtered viral media was added per 15cm^2^ dish (1×15cm^2^ dish per T225 flask), and the cells were incubated for 6 hours at 37°C. Afterwards, the viral media was aspirated, and cells were washed 2X with ice-cold PBS-/-. Cells were then dissociated with 5mL of pre-warmed Trypsin-EDTA for 5 minutes at 37°C, centrifuged at 300g for 4 minutes, and then re-plated in new 15cm^2^ dishes in fresh culture media. Cells were then left to incubate overnight at 37°C. The following day (Day 9) the washing, dissociation, and re-plating steps described above were repeated and cells were placed in a T225 flask in production media. Cells were then left incubating at 35°C. Between Days 10 and 14, 5mL of fresh production media was added to each T225 flask. On Day 14, supernatant media containing pseudotyped RV particles was collected, pooled, filtered using 0.22µm PES vacuum filter, and was incubated with Benzonase Nuclease (1:1000 dilution; Millipore Sigma, Cat no. 70664) for 1h at 37°C. Following incubation, the supernatant was overlaid on a 20% sucrose cushion in an open-top polypropylene tube and placed in a Beckman Coulter Ultracentrifuge in Beckman Coulter SW32Ti rotor. The samples were centrifuged at 20,000 RPM for 2 hours at 4°C. Following centrifugation, the supernatant and sucrose cushion were aspirated from the top and the pellet containing pseudotyped rabies virus was resuspended in 15-30µL of PBS-/-. The virus was aliquoted and stored at −70°C until use.

#### Rabies plasmid and viral library sequencing

To assess barcode diversity of the plasmid library, the barcoded RV plasmid pool was used directly as a template for three nested PCR reactions to amplify the barcode region. PCR1 and PCR2 were performed at an annealing temperature of 53°C, with cycling stopped in the logarithmic phase to minimize amplification bias (20 cycles). In PCR1, 10ng cDNA/reaction was used with primers RV_AMP_PCR1_Primer_1 (5’-CTGAGGGCATGAACTGGGTATACAA-3’) and RV_AMP_PCR1_Primer_2 (5’-GA ACTTCACGTGGTAGATGAGGGTG-3’) to amplify a 900 bp region including the barcode. The PCR product was purified using SPRIselect Beads (Beckman Coulter, Cat no. B23318). PCR2 used staggered primers directly flanking the barcoded region to increase barcode sequencing resolution, using 150 ng template/reaction (7 cycles).

Forward primers included RV_AMP_PCR2_FWD_Primer_1–4: 5’-ACACGACG CTCTTCCGATCTTCTTCTCTGCTTCTAGAATAACTA GGAG-3’, 5’-ACACGACGCTCTTCCGATCTDTCTTCT CTGCTTCTAGAATAACTAGGAG-3’, 5’-ACACGACGC TCTTCCGATCTBDTCTTCTCTGCTTCTAGAATAACT AGGAG-3’, 5’-ACACGACGCTCTTCCGATCTHBDTCT TCTCTGCTTCTAGAATAACTAGGAG-3’.

Reverse primers included RV_AMP_PCR2_REV_Primer_1–4: 5’-GTGTGCTCT TCCGATCTTTGATGTACGCCTTGGAGCC-3’, 5’-GTGT GCTCTTCCGATCTDTTGATGTACGCCTTGGAGCC-3’, 5’-GTGTGCTCTTCCGATCTHDTTGATGTACGCCTTG GAGCC-3’, 5’-GTGTGCTCTTCCGATCTVHDTTGATG TACGCCTTGGAGCC-3’.

The PCR2 product was purified using SPRIselect Beads, followed by an indexing PCR3 reaction (40ng/reaction, 60°C annealing temperature, 12 cycles) with unique NEB dual indexing primers (NEBNext Multiplex Oligos for Illumina (96 Unique Dual Index Primer Pairs), NEB #E6440) and a final SPRIselect purification.

To assess barcode diversity of the viral library, viral RNA was first isolated from the purified RVΔG–UVI–E2-Crimson preparation using the RNeasy Mini Kit (Qiagen, Cat no. 74104), and cDNA was generated using the LunaScript RT SuperMix Kit (New England Biolabs, Cat no. E3010). The same three nested PCR reactions were then performed using the primers described above, with identical cycling parameters and purification steps. For both libraries, amplicons were indexed with unique NEB dual indices, verified for quality and size (540–580 bp) by Fragment Analyzer, and quantified by Kapa qPCR. Libraries were pooled and sequenced at the VBCF NGS facility on an Illumina NovaSeq 6000 S4 flow cell in paired-end 150 bp mode, generating 8.1 million and 8.2 million reads for the plasmid and viral libraries, respectively.

#### Targeted amplification of the UVIs from single-cell libraries

To recover barcode identity from single-cell transcriptomes, we performed targeted amplification of the RV barcode directly from 10X Genomics scRNA-seq cDNA libraries using three nested PCR reactions. In PCR1, primers were designed to amplify a region containing the barcode (RV_AMP_PCR1_Primer_1: 5’-CTGAGGGCATGAACTGGGTATACAA-3’ and RV_scAMP_Read1_REV_Primer: 5’-CCCTACACGACGCTCTTCCGATCT-3’, binding to the 10X TruSeq Read 1 sequence) using 10 ng of purified 10X cDNA library/reaction at 53°C Tm for 15 cycles. The PCR product was purified using SPRIselect Beads (Beckman Coulter, Cat no. B23318). PCR2 used staggered primers directly flanking the barcoded region to increase barcode sequencing resolution, using 75 ng template/reaction at 53°C Tm for 10 cycles.

Forward primers included: RV_scAMP_PCR2_FWD_Primer_1–4: 5’-GTGTGCTCTTCCGATCTTCTTCTCTGCTTCTAGAATAACTAGGAG-3’, 5’-GTGTGCTCTTCCGATCTDTCTT CTCTGCTTCTAGAATAACTAGGAG-3’, 5’-GTGTGCT CTTCCGATCTBDTCTTCTCTGCTTCTAGAATAACTA GGAG-3’, 5’-GTGTGCTCTTCCGATCTHBDTCTTCTCT GCTTCTAGAATAACTAGGAG-3’

The reverse primer was the same TruSeq Read 1 primer used in PCR1 (RV_scAMP_Read1_REV_Primer: 5’-CCCT ACACGACGCTCTTCCGATCT-3’).

The PCR2 product was purified using SPRIselect Beads, followed by an indexing PCR3 reaction (12 cycles, 60°C annealing temperature, 40 ng template/reaction) with unique NEB dual indexing primers (NEBNext Multiplex Oligos for Illumina (96 Unique Dual Index Primer Pairs), NEB #E6440).

#### Transsynaptic tracing by AAV and rabies

The retrograde rabies viral transsynaptic labeling protocol is depicted in Figure 1. Individual 129-day-old organoids were removed from their 10cm dishes and transferred to single wells of a 24-well plate (Eppendorf, Cat no. 0030722116) containing 1.5mL complete BrainPhys media and 5×10^10^ AAV and were incubated overnight. The following day, organoids were removed from the 24-well plate and placed into new 10cm dishes containing 20mL complete BrainPhys media. Organoids were then fed on a normal feeding schedule for 14 days with complete BrainPhys media. 14 days later, the 143-day-old organoids were removed from their dishes and transferred to single wells of a 24-well plate containing 1.5mL complete BrainPhys media and 1×10^5^ viral particles of the RVΔG-UVI-E2-Crimson. Organoids were then placed back at 37°C overnight. The following day, organoids were removed from the 24-well plate and placed back in a 10cm dish containing 20mL complete BrainPhys media. Organoids were placed back on a normal feeding schedule for 7 days until they were 150 days old at which time they were processed for down-stream analyses. Note that our experimental setup for rabies infection is very different from other *in vivo* work. In the mouse brain, tracing is initiated by local injections of highly concentrated rabies virus. In contrast, we use rabies virus at a low concentration, distributed evenly throughout the medium, to allow sparse, single-virion labeling.

#### Viral tracing optimization and validation assays

To evaluate the effect of construct architecture on tracing efficiency, two AAV helper constructs were compared in parallel: a tricistronic construct (CaMKII-TVA^WT^-P2A-N2cG-P2A-EGFP; TVA^WT^-TriCis) expressing TVA, N2cG, and EGFP as separate elements, and a bicistronic fusion construct (CaMKII-TVA^WT^::EGFP-P2A-N2cG; TVA^WT^::EGFP, hereafter TVA^WT^) in which EGFP is fused directly to the TVA locus. The 3^*′*^ regulatory cassette was shortened by replacing the original bGH polyA–WPRE module with the more compact SV40 polyA–WPRE3 module. This removes roughly 440 bp more from the construct, reducing a ~4.9 kb cassette to approximately 4.0 kb, which should bring it back into a more favorable size range for AAV packaging and reduce the risk of inefficient or truncated packaging. Organoids were infected with each AAV construct, followed by RVΔG– UVI–E2-Crimson infection 14 days later. After a further 7 days, organoids were dissociated and the percentage of E2-Crimson^+^ cells was quantified by FACS as a measure of tracing efficiency. To compare tracing efficiency between TVA variants, organoids were infected in parallel with AAVrg carrying either TVA^WT^ or TVA^66T^ (CaMKII-TVA^66T^::EGFP-P2A-N2cG), using the same 14-day AAV and 7-day RV intervals. The percentage of E2-Crimson^+^ cells was compared between conditions to select the optimal TVA variant for subsequent experiments. To distinguish transsynaptic spread from nonspecific RVΔG uptake and to quantify spread rate, a glycoprotein knockout control (CaMKII-TVA66T::EGFP-P2A-N2cG^M1^*,^P440^*; TVA^66T^(GKo)) was generated by introducing dual stop codon mutations at residues M1 and P440 of N2cG, abolishing glycoprotein function while preserving TVA expression and starter cell formation. An equivalent GKo construct was generated for TVA^WT.^ Organoids were infected in parallel with either the functional construct or its matched GKo control for both TVA variants, followed by RVΔG–UVI–E2-Crimson infection 14 days later. Organoids were dissociated and analyzed by FACS 7 days post-rabies infection. Because GKo organoids support starter cell formation but not transsynaptic spread, the mean percentage of E2-Crimson^+^ cells in the GKo condition provided a specific estimate of the starter cell baseline. Spread rate was calculated by dividing the mean percentage of E2-Crimson^+^ cells in the functional construct condition by the mean percentage in the matched GKo condition, enabling a comparison of spread rates between TVA^WT^ and TVA^66T^.

#### Dual-color co-infection assay / single virion infection (UVI) per cell

The scRNA-seq data do not provide a ground-truth estimate of true multi-virion infection. A multi-UVI call in scRNAseq, in an AAV+ rabies+ cell can reflect true infection by multiple UVI-bearing rabies virions, but can also arise from residual ambient UVI RNA, incomplete barcode correction, cell doublets, or differences in cellular or RNA recovery. We therefore estimated the frequency of direct multi-virion infection using two-color control experiments with separate RVΔG-EGFP and RVΔG-E2-Crimson preparations followed by FACS analysis.

To obtain an upper estimate, we performed these experiments in HEK293T cells stably expressing TVA (HEK-TVA, gift from ISTA) ^96^. Therein, all cells are accessible to infection, and express high levels of genomically integrated TVA. These conditions are not guaranteed and could vary in AAV-transduced organoids. HEK-TVA cells were grown in 6-well plates to 90–100% confluency and co-infected with two RVΔG preparations carrying distinct fluorophores at a total of 2 × 10^5^ (**Fig. S1H**) or at 4 × 10^5^ virions per well, split equally between GFP and E2-Crimson viruses. These concentrations corresponded to 2–4 times higher virion counts than what we have used in tracing conditions. The raw observed mixed-color dual-positive rates were 0.03% and 0.01%, respectively.

To account for same-color co-infections not directly detected by the two-color readout, we modeled infection as a Poisson process. The observed GFP+ and E2-Crimson+ fractions, pGFP and pCrimson, were converted into inferred Poisson parameters as λGFP = −ln(1 − pGFP) and λCrimson = −ln(1 − pCrimson). The parameters were summed as λtotal = λGFP + λCrimson. The expected fraction of cells receiving two or more infection events was then calculated as P(k≥ 2) = 1 − exp(−λtotal) × (1 + λtotal), yielding corrected multi-virion infection rates of 0.065% and 0.018% of all cells, respectively.

#### Dissociation of organoids

Individual organoids were transferred directly into gentleMACS dissociation tubes (Miltenyi Biotec, Cat no. 130-096-334) containing 2 mL of a 1:1 Trypsin-Accutase solution supplemented with DNase I (Thermo Fisher Scientific, Cat no. 89836) at a final concentration of 50 U/mL (100 U per organoid). Tubes were run on the gentleMACS dissociator using the 37C_NTDK_2 program (~29 minutes). Following dissociation, samples were diluted with 13 mL of room-temperature PBS supple-mented with 0.04% BSA (Sigma-Aldrich, Cat no. A8806-5G). The cell suspension was then passed dropwise through a pre-wetted 70 µm cell strainer (Miltenyi Biotec, Cat no. 130-098-462) into a 15 mL Falcon tube (Corning, Cat no. 352095). Samples were centrifuged at 300g for 10 minutes at 4°C to pellet cells. The supernatant was aspirated, leaving approximately 20 µL of residual PBS, and the pellet was resuspended in 800 µL PBS supplemented with 0.04% BSA, Calcein Blue (Thermo Fisher Scientific, Cat no. C1429; 5 µM working concentration), and DNase I (20 U). The suspension was passed through the 35 µm filter of a FACS tube that had been pre-wetted with PBS, and samples were kept on ice until sorting or acquisition.

#### Immunofluorescent staining

Individual organoids were removed from media and placed in wells of a 24-well plate (Eppendorf, Cat no. 0030722116) with 1mL of 4% PFA (Sigma-Aldrich, Cat no. 441244) diluted in Dulbecco’s phosphate-buffered saline (DPBS) (Gibco, Cat no. 14040174) and fixed for 90 minutes with gentle shaking at room temperature. After fixation, 4% PFA was removed and replaced with 1mL DPBS. Organoids were then shaken gently for 10 minutes. This was repeated 3 times. Following these washing steps, DPBS was removed and replaced with 2mL 30% sucrose, diluted in DPBS. 24-well plates were then sealed with parafilm, and organoids were transferred to 4°C where they were kept gently shaking overnight or until they sank to the bottom of the well. Once the organoids had sunk to the bottom of the wells, they were then embedded in O.C.T. (Sakura Finetek USA Inc., Cat no. 4583). This involved first removing 1mL of the sucrose solution and replacing it with 1mL O.C.T., creating a 50:50 mixture. The solution was then mixed thoroughly with a pipette tip making sure not to damage the organoids. The organoids were then placed on a shaker and gently mixed for 60 minutes at room temperature. After the 60-minute incubation, the 50:50 mixture was removed and replaced with 1mL 100% O.C.T. Organoids were then removed from the wells with a p1000 pipette tip that was cut to create a wide bore to allow the entire organoid to be removed. The organoids were then transferred carefully into Disposable Base Molds (Electron Microscopy Sciences, Cat no. 62352-15), filled with O.C.T. and placed on top of an aluminum block that had been cooling in dry ice, making sure it was flat atop the dry ice to avoid shifting of the organoids. Organoids were then allowed to freeze in the plastic molds and stored at −70°C. Cryoblocks were sectioned at 20 µm thickness using a cryostat (Thermo Fisher Scientific, CryoStar NX70) and stored at −20°C for long-term storage and downstream immunostaining which was performed as follows. Organoid sections were thawed and allowed to dry at room temperature. A liquid blocker pap-pen (Merck, Cat no. Z377821) was used to draw a border on the perimeter of the slide to prevent liquid from spilling over the slide. The border was allowed to dry at room temperature. Slides were then washed 3 times for 5 minutes in DPBS. Slides were then permeabilized and blocked by incubation in blocking solution (0.25% Triton X-100 (Sigma-Aldrich, Cat No. 93420), 4% Normal Donkey Serum (EMD Millipore, Cat no. S30-100ml) diluted in DPBS) for 60 minutes at room temperature in a humidified chamber. Antibody incubations were performed in antibody solution (0.1% Triton-X, 4% Normal Donkey Serum in DPBS). First, sections were incubated with antibody solutions containing diluted antibodies overnight. The next day, slides were washed 2x for 5 min with PBS-T (0.05% Triton-X in DPBS) and 1x for 5 minutes with DPBS. Sections were then incubated with an antibody solution containing secondary antibodies [1:1000] and Hoechst 33342 nuclear dye for 2 hours at room temperature. After secondary antibody incubation, sections were washed 3 times for 5 minutes in DPBS and mounted with coverglass using DAKO mounting medium. Slides were left at room temperature to dry overnight while protected from light and kept at 4°C for long-term storage.

#### Image acquisition

Immunostaining images were acquired with an SP8 confocal microscope (Leica) and a Panoramic FLASH 250 II digital Slide Scanner (3DHISTEC) at 10 and 20x magnification. Live imaging was done using an Olympus SpinSR10 spinning-disc microscope.

#### Calcium imaging recordings and analysis

TSC patient and isogenic control iPSC lines were genetically engineered by CRISPR/Cas9 to stably express GCaMP6s under the human synapsin promoter (hSyn1-GCaMP6s); integrated at the AAVS1 locus. Organoids were generated as described above (Methods; Organoid Generation) and imaged at 150 DIV on an Olympus IX83 inverted microscope with a Yokogawa W1 dual-camera spinning disk, a 20x objective, and Hamamatsu Orca Flash 4.0 cameras (×3.2 additional magnification). Recordings were acquired at 15.4 frames per second for 5 minutes and 25 seconds in BrainPhys medium under controlled environmental conditions (~37 °C and 5% CO_2_). Samples were pseudonymized before acquisition, and all analysis was performed blind to genotype. Recordings were downsampled to 512 × 512 pixels and the first 100 frames were removed. Fluorescence traces were extracted with the open-source software package CaImAn (github.com/flatironinstitute/CaImAn) ^98^, filtered to remove artifacts, and z-scored. Recordings with fewer than three functional neurons were excluded. Only batches with matched control and mutant pairs at the same time point were included (Controls: 26 organoids, 57 recordings; TSC: 24 organoids, 57 recordings; 114 recordings in total, **Table S3**).

Calcium network events were detected on the population-averaged z-scored trace above 1.5 x baseline standard deviation, estimated by median absolute deviation (MAD). Network events were defined as contiguous segments with an internal peak ≥ 3 × baseline s.d. and duration > 0.5 s. Network events separated by < 0.3 s were merged. For each network event, we computed frequency, participation fraction, event duration, decay time, rise time, and event amplitude, and counted participating neurons (max z-score > 1.0 within the event window) on events with > 3 participating neurons. All analysis was implemented in Python with custom code that will be available at github.com/knoblichlab/org_calcium_imaging.

#### FACS analysis and sorting

For flow cytometry analysis, the NovoCyte Penteon Flow Cytometer was used. For cellular isolation for downstream scRNA-seq, a BD FACS Aria III was used with an Alexa 700 filter at 20 PSI (100µm nozzle). Cells were collected in DPBS supplemented with 4% BSA. For each sample, all E2-Crimson^+^ cells were collected. Cells from the E2-Crimson^-^ population, equivalent to ~10% of the total E2-Crimson^+^ cell count were also collected for sequencing to establish a snapshot of the organoid’s cell type composition for downstream analyses (CTC). For each experiment, cells from an organoid of a distinct genetic background that had not been exposed to AAV or RVΔG–UVI– E2-Crimson were sorted separately and mixed into the single-cell solution at a minimum of 5% or 1000 cells, enabling the estimation of library-specific ambient viral RNA contamination.

For sorted samples, prior to library preparation, FACS-sorted fractions were subjected to a gentle centrifugation step to concentrate cells and remove excess sort buffer. Samples were centrifuged at 400g for 5 minutes at 4°C. The expected recovery volume was calculated to target a final concentration of 450–1000 cells/µL. The supernatant was carefully removed in 40 µL increments using a wide-bore pipette tip (Thermo Fisher Scientific, Cat no. 2069GPK), using a reference tube to gauge the remaining volume, until the calculated target volume was reached. Removed supernatant was retained in a separate tube and checked to assess cell loss from this step using a NC-250 cell counter (Chemometec, Product no. 970-0251). The remaining volume was confirmed, and the pellet was gently resuspended using a wide-bore pipette tip. Resuspended cells were kept on ice and immediately transferred for scRNA-seq preparation and sequencing.

#### Nested multiplexing

To enable joint sequencing of traced and control cells from the same organoid, dissociated tracing organoids were separated by FACS into an E2-Crimson-positive traced fraction and an EGFP-negative/E2-Crimson-negative cell type composition control (CTC) fraction. All traced cells and a small fraction of CTC cells were collected. In parallel, live singlets were isolated from a typically age-matched organoid of a genotypically distinct cell line that had not been exposed to AAV or RV, and served as true-negative (TN) cells. Traced, CTC, and TN cells were pooled into a single 10x Genomics library. After sequencing, all three populations were computationally separated, as described under “Nested demultiplexing to identify RV+, AAV+ (and UVI+) cell populations”.

#### Single-cell RNA-seq library preparation

For each library, cells were counted and loaded onto a 10x Chromium controller. Libraries using the Chromium Single Cell 3’ Reagent Kits (v.3.1, Cat No. PN-1000121) were prepared following the 10X user guide. Libraries were sequenced on a NovaSeq S2 or S4 flow cell with a target of 50,000 paired-end reads per cell.

#### Whole Genome Sequencing (WGS)

For WGS of cell line published in this study, we followed the same protocol as for the published lines under EGAD00001008769.

## Computational Methods

### Single-cell Analysis

As in ^48^, we first aligned reads to the GRCh38-3.0.0 human reference genome, but extended with the rabies and AAV constructs, using Cell Ranger v6.0.1 (10x Genomics) with pre-mRNA gene models and default parameters. This produced a cell-by-gene, umi count matrix. UMI counts were then analyzed in R v4.4.1 with a Bioconductor 3.19 package container, and Seurat v5.2.1^47^. We filtered the cell-by-gene matrix for high quality cells based on the number of genes detected, with a hard minimum of 750 genes, and a dynamic, per-library upper cutoff at the 0.9975 gene quantile, removing cells with high mitochondrial [20%] or ribosomal [40%] messenger RNA content, using the Seurat.utils R package.

Next, we followed the standard Seurat pipeline per library with the following details: count matrices were log-normalized and scaled to 10,000 UMIs per cell, with “library” and “nuclei_class” (intact cells or nuclear fragments) regressed out; expression of the 2,000 most variable features was Z-scaled; and principal component analysis (PCA) was performed. We imported UVI annotation per library and processed it as described under “UVI filtering and processing”. Downstream steps were performed for data inspection. The data presented in the paper were re-analyzed together after individual datasets had been filtered, curated, and merged (below).

#### Nuclei classification

As sensitive live neural cells are often damaged during 10x Next GEM encapsulation, some cells partially lose their cytoplasm, while retaining an intact nucleus. These still contained high levels of above-ambient rabies reads and UVIs, and yielded high-quality transcriptomes, but their transcript-class distributions differed from those of whole cells. To account for this latent structure in the data, we classified nuclei-enriched and whole-cell transcriptomes and used this annotation for downstream regression. For each single-cell library, we quantified the nuclear fraction of transcripts using DropletQC ^99^. Cells with nuclear fraction below 0.15 were classified as “cytoplasmic fragments”, and cells with fewer than 1,500 detected genes were classified as “low-quality fragments”. Remaining cells were reclassified as “whole cells” or “nuclei” using unsupervised clustering in the joint space of nuclear fraction and detected gene number. Both k-means clustering and Gaussian mixture modeling were evaluated after feature scaling. The cluster with higher median nuclear fraction was annotated as nuclei and the lower-fraction cluster as whole cells. Although both approaches gave similar results, the k-means clustering performed more reliably and its annotation was used for downstream analyses.

#### Analysis of whole genome sequencing data, cell line genotype demultiplexing and genotypic doublet detection

Cell-line WGS samples were processed using the nf-core/Sarek pipeline v2.5.1 with GATK v4.1.2.0 Haplotype-Caller ^100^ using reference files from the GATK GRCh38 iGenomes bundle. In each 10x library we mixed 2–4 cell lines with different genomes. We therefore performed genotype-based demultiplexing using souporcell ^50^. The pipeline was run with default settings, providing the pre-determined genotype variants through the *known_genotypes* parameter, the Cell Ranger .bam, the Cell Ranger filtered barcodes file, and the reference .fasta as input. This classified cells into donor-genotype singlets, unassigned cells, and genotype doublets. Here, genotype doublets represent cells with multiple donor genotypes and therefore correspond to heterotypic doublets. Unassigned cells were low-quality cells with distinctly low nFeature and nUMI values. For all downstream analyses, all unassigned cells and all genotype doublets were excluded. Following genotype demultiplexing, each pooled library was further partitioned into biological samples by combining donor-genotype assignments with the nested multiplexing strategy described below, which separated rabies-traced and cell-type-control fractions within genotype-matched pools.

#### scRNA dataset integration

The combined scRNA-seq dataset was integrated using Harmony ^51^ on the first 50 principal components. Batch correction used 10x library and nuclei-class assignments. The resulting 50-dimensional Harmony embedding was used for all downstream steps: shared nearest-neighbor graph construction, clustering, and UMAP visualization. Integration was evaluated visually by library and annotation mixing. Downstream processing followed the standard Seurat workflow. Highly variable genes were identified, expression values were scaled, shared nearest-neighbor graph construction, UMAP, graph-based clustering, and DGEA were performed. For the basis of cell type annotation (v11.fine) we used clustering resolution 0.6.

#### Annotation of stressed cells using gruffi

Stressed cells were identified using the gruffi framework ^48^, which detects stress-associated transcriptional programs by partitioning the data into hundreds of small transcriptomic clusters (“granules”) of approximately 100 cells each. For each granule, average expression scores were computed for predefined gene sets representing stress-associated and non-stress programs. Stress signatures included glycolytic process (GO:0006096) and endoplasmic reticulum stress (GO:0034976), while gliogenesis (GO:0042063) and astrocyte differentiation (GO:0048708) were used as non-stress reference programs to distinguish physiological, lineage-associated transcription from stress-induced states. Granule-level score distributions were used to identify clusters with elevated stress program activity in a data-driven manner, avoiding fixed cutoff selection. Stressed cells, together with mispatterned cells, were subsequently excluded from down-stream analyses.

#### Exclusion of mispatterned and stressed cells

Following stress annotation, mispatterned cells were identified by transfer of coarse patterning labels from a comprehensive organoid reference dataset ^49^ to the connectomics Seurat object, using label transfer (FindTransferAnchors and TransferData). Cells assigned to stromal, eye, oligo, choroid plexus, or floorplate progenitor identities were classified as mispatterned and excluded. Small RNA_snn_res.0.2 clusters 8 and 11 were additionally excluded because they (i) separated distinctly in UMAP space, (ii) were almost completely dominated by mispatterned and (iii) stressed cells, indicating artefactual and non-neural states. In total, ~2.3% of cells were excluded as stressed, ~3.3% as mispatterned, ~1.6% as both stressed and mispatterned, and ~0.6% by complete exclusion of clusters 8 and 11, leaving ~92% of cells for downstream analyses.

#### Quantification of (anti)viral response across cell states

To assess whether viral infection or the associated host response affected the experimental readouts, we quantified per-cell gene-set scores for antiviral innate immune response (GO:0140374), response to type I interferon (GO:0034340), canonical NF-κB signal transduction (GO:0007249), and defense response to virus (GO:0051607) across cell states. Scores were summarized by calculating the median for each cell state and were additionally evaluated by visual inspection. We also examined the viral response genes reported by Lee et al., 2023^54^. Together, these analyses indicated that the antiviral response remained localized rather than broadly affecting cell states, consistent with the absence of resident immune cells in the organoids.

#### Seurat clustering and final cell-type annotation

We performed shared nearest-neighbor (SNN) clustering in Seurat. A k-nearest-neighbor graph was constructed from Euclidean distances in 50-dimensional PCA space, and SNN edge weights were calculated using Jaccard similarity between shared neighbors. Louvain clustering was performed at resolutions 0.3 and 0.6. The resolution 0.6 solution was used for fine cell-state annotation. Cell-cycle scores were calculated with Seurat’s “CellCycleScoring” function and default parameters. Cycling progenitors in G2M and S phase (clusters 10 and 12) were merged into a single “Prog, Dividing” state; cell-cycle effects were not regressed out at this stage.

Radial glia (RG) and astrocytes were not resolved by transcriptome-wide clustering. We therefore subsetted this population and reclustered it using the same workflow after regressing recalculated cell-cycle scores. Reclustering at resolution 0.1 identified five clusters. Astrocyte identity was scored with Seurat’s “AddModuleScore” using *S100B, GFAP, ALDH1L1, AQP4, CD43, CHI3L1, APOE, BHLHE41, MOXD1, FAM107A, TNC, ANXA1, CD38, CLU*. Clusters with positive median module scores were assigned as astrocytes; clusters with negative scores were assigned as RG. Marker expression was inspected as an additional validation.

We provide three additional levels of annotation: “Cell.types.v12.coarse” (based on resolution 0.3, 12 cell types, RG/astrocyte separated as above), “Lineage.ExN” (Progenitor; IN; Glia; ExN, UL; ExN, DL) and “Cell.Class” (Neuron; Progenitor; Glia). Broader annotation levels were generated by merging categories of the fine cell-state annotation.

#### Label transfer from cortical datasets and cluster annotation

For cluster annotation, we performed reference-based label transfer from our earlier work ^49,53^as well as fetal data (cellxgene: ad2149fc-19c5-41de-8cfe-44710fbada73) ^52^, using Seurat’s implementation with default parameters. Final cluster labels were assigned by comparing transferred annotations with marker gene expression (Figure 2C, S2A). With the fully annotated and preprocessed data, we then analyzed the joint connectivity and gene expression data.

To validate the *in vivo* identity of the upper-layer excitatory neuron clusters, we performed reference-based label transfer from additional human fetal brain datasets using SingleR ^101^, subsetting the temporal windows to broadly align with the developmental stages of our organoids. We used the following public *in vivo* reference datasets for label transfer: (1) GW15–22 (H1) data from Zenodo:14422018 and merscope_integrated_855.h5ad ^59^; (2) GW22–24 data from https://brain.listerlab.org, under Processed_data, “hsBrH_ga22.24_majorcl”, corresponding to (GSM5138509 ga22_snRNAseq; https://www.ncbi.nlm.nih.gov/geo/query/acc.cgi?acc=GSM5138511GSM5138511 ga24_snRNAseq)^60^; (3) GW17–18 data from http://solo.bmap.ucla.edu/shiny/webapp [corresponding to dbGaP phs001836.v1.p1]^61^. At late gestation, only the latest born UL-ExNs are expected to migrate; thus Migrating ExN cell type annotations ^59,61^, were counted towards upper layer ExNs. For the label transfer, only reference clusters with more than 50 cells were kept, and reference profiles were aggregated into pseudo-bulk cluster medians (aggr.ref = TRUE). Feature selection across reference groups was driven by pairwise Wilcoxon rank-sum differential expression tests (de.method = “wilcox”), retaining the top 20 marker genes per pair (de.n = 20) to compute rank correlation assignments for query cells. All other parameters were left default.

#### Nested demultiplexing to identify RV+, AAV+ (and UVI+) cell populations

In scRNA-seq, closely related cell populations such as experimental subsets and internal controls are often best sequenced in the same library to reduce cost, handling time, technical variability, and cell loss. This creates a general demultiplexing problem when the target populations are biologically similar, share the same genotype, or cannot be cleanly separated by endogenous marker expression after sequencing. Directly labeling the target populations can solve this problem, but additional labeling steps may be incompatible with fragile, rare, or time-sensitive samples. This is especially problematic in lineage tracing, synaptic network mapping, or in rare cell type experiments, where cell loss and delayed library preparation can compromise data quality. NestedMultiplexer addresses this problem by adding a third, ground-truth negative population to the same library. This negative population does not need to be precious experimental material and can therefore be marked independently, for example by genotype, antibody- or lipid-based cell hashing, or any orthogonal identifier. After sequencing, this externally identifiable negative population provides a library-specific baseline for ambient marker RNA and technical background. The two target populations can then be separated using an experiment-specific threshold (e.g. FPR=0.01), corresponding to a cutoff-value at which only 1% of true-negative cells are classified as positive.

Nested demultiplexing was applied at three distinct stages of our pipeline. First, true-negative control cells provided a library-specific baseline of ambient viral RNA, enabling the precise thresholding and classification of rabies-traced cells versus uninfected cell-type composition control (CTC) cells. Second, true-negative cells enabled parameter optimization to establish library-specific optimal read- and UVI-depth cut-offs for identifying bona fide unique viral identifiers (UVIs) while minimizing false-positive assignments (described under “Optimal Read and UVI-umi threshold identification”). Finally, nested demultiplexing facilitated the robust identification of starter cells based on the detection of AAV-derived transcripts.

In our experiments, each 10x library contained three cell populations: (1) traced cells from traced organoids, sorted as E2-Crimson-positive (*Trace*); (2) cell type composition control cells (*CTC*) from the *same* organoids, sorted as EGFP- and E2-Crimson-negative; and (3) true-negative (*TN*) cells from an unrelated, never infected, typically age-matched organoid from a different genotype. Because Trace and CTC cells came from the same organoid, they shared the same genotype and could not be separated by genotype demultiplexing alone. In contrast, TN cells were expendable control material, from typically age-matched organoids of a different genotype never exposed to virus. This allowed their unambiguous identification, and TN samples were then used to estimate the library-specific background of ambient rabies RNA. This control was necessary because low levels of rabies-derived transcripts were detectable in all cell fractions, including rabies-negative cells, due to ambient RNA released during dissociation.

We therefore first used genotype demultiplexing to identify the true-negative cells. True-negative cells were defined as high-quality singlets assigned by *souporcell* to the uninfected donor genotype. For each cell, total rabies expression was calculated as the sum of normalized expression values of the rabies-derived transcripts N, P, M, E2-Crimson, and L from the RNA assay log-normalized data layer. Within each library, we then calculated the 95th percentile of the log10-transformed total rabies expression across true-negative cells and used this value as the library-specific threshold to distinguish rabies-positive *Trace* cells from rabies-negative *CTC* cells. This avoided arbitrary global cutoffs and accounted for library-to-library differences in ambient rabies RNA contamination. The only exception was library sc15.033, where TN samples were of low quality, therefore, after thorough inspection, we set the threshold to the 90^th^ percentile.

#### Quantification of rabies-associated transcriptomic mixing using LISI

To determine whether rabies tracing altered transcriptomic state, we quantified the local intermixing of CTC (RV-) and Trace (RV+) cells within each annotated cell state. Analyses were performed within cell states to control for differences in cell-state abundance between conditions. Local mixing was measured using the Local Inverse Simpson’s Index, LISI ^51^. For each cell, LISI constructs a distance-weighted neighborhood in the selected embedding, such that nearby cells contribute more strongly than distant cells. The weighted proportions of CTC and Trace cells are squared and summed, and the reciprocal of this sum is reported as the LISI score. For two balanced populations, a score of 1 indicates a neighborhood containing only CTC or only Trace cells, whereas a score of 2 indicates equal representation of both populations. Higher values therefore indicate stronger local intermixing. Cell states containing at least 100 cells in each condition were retained. Within each cell state, CTC and Trace populations were balanced by random downsampling to the size of the smaller population. This procedure was repeated 5 times to assess the stability of the estimated mixing scores. LISI was calculated with perplexity 30 and nearest-neighbor tolerance nn_eps = 0.1. Calculations were performed in the existing PCA, Harmony, and UMAP embeddings. The first 50 dimensions were used for PCA and Harmony, whereas both UMAP dimensions were used for the UMAP analysis. The PCA embedding represents the concatenated dataset before integration. Harmony integration corrected for Library and “nuclei_class”. Rabies infection or tracing status was not included as a covariate in either Harmony integration or data scaling. For each cell state and downsampling replicate, the median cell-level LISI score was calculated. Replicate medians were then averaged within each cell state. The reported value for each embedding was calculated as the mean of these cell-state-specific median LISI values. Variation across cell states was summarized using the coefficient of variation, calculated as the standard deviation of the cell-state-specific values divided by their mean.

#### Rabies-associated transcriptomic shifts relative to biological variation

To quantify rabies-associated transcriptional perturbation independently of cell-state composition, RV-positive Trace and RV-negative CTC cells were compared within each annotated cell state, and True Negative cells were excluded. Comparisons were restricted to groups containing at least 100 cells: both Trace and CTC populations for within-state rabies comparisons, each CTC population for pairwise cell-state comparisons, and each pooled Trace and CTC cell-line population for within-state cell-line comparisons. The analysis was restricted to coding host genes with non-zero expression in at least 1% of the cells. Viral genes were removed. Gene counts were log-normalized in Seurat, and gene-wise log2 fold changes were calculated using the same gene universe for every comparison. Each comparison was summarized by the root mean square of its gene-wise log2 fold changes. This metric captures the overall magnitude of transcriptomic separation irrespective of direction, with larger values indicating broader or stronger expression differences. The rabies-associated effect was calculated from Trace-versus-CTC comparisons within each cell state. Typical cell-state variation was defined as the median effect across pairwise comparisons of each CTC state compared with all other eligible CTC states. Typical cell-line variation was defined as the median effect across pairwise cell-line comparisons within the same cell state, pooling Trace and CTC cells within each line to improve statistical power. Relative effects were then computed by dividing the rabies-associated effect by each reference separately; values below 1 indicate that the rabies-associated shift is smaller than the corresponding cell-state or cell-line variation. These ratios were used descriptively to compare effect magnitudes, without formal statistical testing.

### UVIs

#### UVI-identification in bulk

For UVI-calling in bulk libraries, we first counted then discarded uncut sequences (no UVI between flanks). We then used Cutadapt v1.8^102^, to identify reads with correct 5’ and 3’ UVI-flanks (10% error tolerance, min 6nt match), trimmed those flanks away, and selected exact 14-mers from the remaining sequences. We filtered for the synthesis pattern: [ACT][ACGT][CGT][ACGT] [ACG] [ACGT][AGT][ACGT][ACT][ACGT][CGT][ACGT][ACG] [ACGT]. Finally, sequence correction was performed using Shepherd ^45^, with default settings, on all UVIs. We then counted reads for each distinct UVI sequence.

#### UVI-identification in single-cell libraries

Sequences were aligned using Cell Ranger v6.0.1 against a reference genome extended with the viral constructs. Following alignment, the TAMP and GEX alignments were merged using Samtools (v1.15.1) ^103^. To focus on the relevant sequences, reads mapping to the “M_UVI.RabV” sequence were selected and subsequently converted to .fastq format with Samtools (v1.15.1).

Preprocessing steps followed established protocols similar to those used in bulk experiments. First, Cutadapt v1.8 was used to identify reads containing correct 5’ and 3’ UVI-flanking sequences, allowing for a 10% error tolerance and requiring a minimum of 6 nucleotide matches. These flanking regions were then trimmed away, leaving only exact 14-mers for further analysis. Sequence correction was performed using Shepherd ^45^, with default settings, on all UVIs. This resulted in an average 85% reduction of detected raw barcodes. Next, we selected reads originating from bona fide cell barcodes (CBCs) based on the read IDs in the .bam files. UVIs matching known synthesis patterns were selected, and raw, unfiltered count tables were generated and stored in the subfolder “UVI.”

#### UVI library comparison across publications and representation bias quantification (bulk)

Barcode/UVI abundance distributions were compared between published rabies barcode libraries and the RVΔG–UVI–E2-Crimson library generated in this study. Source count tables were converted to barcode-level abundance vectors where required. To reduce preprocessing-dependent differences between datasets, barcodes supported by two or fewer reads were first removed from all libraries, as published count tables differed in low-read-count filtering and barcode-correction procedures, and low-read-count UVIs contributed disproportionately to these differences. Remaining barcode counts were ranked by decreasing abundance and renormalized within each library as the percentage of total retained barcode reads. For the RVΔG–UVI–E2-Crimson library, sequence-corrected UVI counts were used. These ranked percentage distributions were used to generate full rank-frequency curves, top-25 barcode frequency plots, and cumulative rank-frequency curves across libraries.

For library bias quantification, we developed a simple metric, the 90% width of the library (q90WL), to quantify complexity and representation bias in barcoded rabies libraries. The 90% width is defined as the minimum number of UVIs required to account for 90% of all sequencing reads in the library, or, equivalently, the number of most frequent (ranked) barcodes that together contribute 90% of the reads. This provides a tangible and intuitive measure of the lower bound of “useful” library complexity: it approximates the minimum number of distinct barcodes that one is likely to observe in downstream experiments. A key advantage of this metric is its interpretability as a concrete count, in contrast to abstract indices such as the entropy metrics. Because the 90% width is determined by the most frequent barcodes, it has two more key advantages: (1) it is largely unaffected by sequencing depth within the typical range of experiments, as the frequencies of top barcodes are reliably estimated even from relatively few reads (unlike Shannon entropy or richness); and (2) it is insensitive to often large differences in rare barcode counts that result from different preprocessing choices, or error-correction strategies. At the same time, it directly reflects representation bias within the abundant fraction of the library. The q90WL metric also has limitations: it always strongly underestimates the real experimental barcode complexity, because of its definition emphasizing robustness. It becomes unreliable at low sequencing depths and when libraries have undergone substantially different preprocessing. For complementary assessment, we also computed Shannon entropy of the barcode distribution, which summarizes the overall evenness of the library. Together, the 90% width and entropy capture both the practical and theoretical diversity of barcoded libraries (Figure S1L–M).

#### Pairwise Hamming distance calculation across rabies UVIs

To estimate the risk that sequencing or PCR errors could convert one valid UVI into another valid UVI during barcode assignment, we estimated the pairwise sequence separation across UVIs. To assess sequence separation within the corrected rabies 14-bp UVI library, pairwise Hamming distances were calculated among distinct UVI sequences from bulk barcode sequencing after Shepherd-based sequence correction. For this analysis, 5,000 unique UVIs were randomly sampled without replacement. All-against-all Hamming distances, excluding self-comparisons were then computed using the stringdistmatrix function with the Hamming metric. The resulting distribution of pairwise edit distances was summarized and visualized as absolute counts and percentages of all non-self pairwise comparisons. The lowest-distance tail was plotted separately to quantify rare barcode pairs with minimal sequence divergence on an absolute scale.

### UVI filtering and processing

#### Optimal read and UVI-umi threshold identification and evaluation of targeted amplification libraries

We have per-formed targeted amplification for the UVIs (see: Targeted amplification of the UVIs from single-cell libraries) and found that targeted amplification increased both the signal (UVI recovery among RV+ cells), and noise (false-positive UVI identification in TN). Therefore, UVI tables including (“combined”) or excluding (“onlyGEX”) targeted amplification were processed independently throughout the following steps.

#### (F1) Library-specific UVI threshold optimization and true-negative validation

Ambient RNA contamination, UVI recovery and other parameters varied across sequencing libraries; thus, simple global thresholding is insufficient for optimal UVI identification. Therefore, using true negative and RV+ target cells, we tested all combinations of different minimum-read (R-)thresholds (T1, T2, T3, T4, T5, T10), requiring min. 2, 3, 4, 5, 6, or 11 reads per UVI-UMI in a cell, respectively, for the UVI to be included. Therefore, we tested each combination of minimum read- and UVI-umi thresholds on the raw UVI tables, separately for GEX-only and combined GEX+targeted amplification UVI calls. We then designed 2 paradigms to find the optimal filtering parameters within the above scenarios for each library. This optimization was possible because of our nested-multiplexing experimental design, which included a true-negative sample in each library for validation. UVI identification in these cells should theoretically be 0%. In the parameter search, we aimed to (a) maximize the UVI recovery rate (>0 UVI) in the maximum number of rabies positive cells (true positive rate) while minimizing either (**b1**) recovery in true negative cells (false positive rate^TN^), or (**b2**) recovery of multiple UVIs in rabies positive cells (proxy false positive rate^R**V**^). For each library, we separately determined the optimal [R,U] parameter set for **a-b1** and **a-b2** scenarios, using an objective function that weighted the true-positive-rate relative to the false-positive penalty ratio of 2:1. We have found that the identified thresholds were largely identical, and that the **a-b2** scenario is always optimizing **a-b1**. We therefore used the **a-b2** optimum read- and UVI-umi-thresholds separately for each library and then validated each by checking the UVI identification rate in true negative cells (b1). In (**Figure S2I**), we compared thresholds resulting from UVI-threshold optimization by the a-b1 or a-b2 strategies and showed higher stringency in b2. Note that we recommend the b1 scenario for *in vivo* applications which use high-concentration local rabies injections where multi-virion infections are common. False positive based optimization is only possible in sparse infection protocols which experimentally show that multi-virion infections are negligible. We implemented library-level, algorithmic decisions on the inclusion of UVI targeted amplification data. Therein, we compared the two optimized solutions (“onlyGEX” and “combined”) using the same recovery versus multiplicity objective and 2:1 weighting. We extracted the recovery and multiplicity rates. We plotted the difference between the “onlyGEX” and “combined” solutions and used the same 2:1 “recovery versus multiplicity” objective function to select the preferred UVI source for each library. Combined GEX+targeted amplification data were se-lected for 4/27 libraries. The selected F1 UVI calls were then passed to downstream steps. **(F2) Low frequency UVI filter**. To suppress residual low-frequency barcodes, if there was at least a fivefold difference between the UMI counts of the top and an additional barcode in a cell, those were discarded. These likely represent residual contamination, and in any case, due to their low cellular presence would lead to a negligible amount of tracing. **(F3) Multiplicity and transcriptomic doublet filter**. Among cells carrying more than one UVI after F2, those independently classified as transcriptomic doublets by scDblFinder using default parameters ^104^, were excluded (genotypic doublets were already excluded earlier). Remaining cells with three or more retained UVIs were also removed from the analysis. **(F4) The library bias filter – Remove cells with non-unique UVIs**. As viral libraries do not have a perfectly even barcode distribution at infinite complexity, multiple cells can receive the same UVI, and thus appear as a single network. To account for this: (i)We measured our libraries complexity and bias by targeted bulk viral-RNA sequencing (**Figure 1 and S1**). (ii) For each sample, we counted distinct UVIs detected in at least two cells after F3 filtering. This *observed* network count can underestimate the number of initially infected cells because independent infections can carry the same UVI. (iii) Therefore, based on the ground truth UVI distribution from the bulk-sequenced rabies library, and the observed number of starter cells, we estimated the expected number of initial infections under the sampling model, which would be observed with a perfectly unique and unbiased rabies library. (iv) Finally based on the estimated initial infection count (number of starter cells), we determined a threshold for the set of barcodes too frequent for the given experiment. Instead of using the lenient threshold of expected value, we opted for a stricter criterion requiring that any UVI should have less than a 5% chance to occur more than once in the given experiment. These probabilities were calculated by simulated sampling of the true number of starter cells from the ground truth rabies barcode library distribution. Barcodes that did not fit the criteria were discarded. **(F5) Resolution or removal of residual multi-UVI cells**. After F4, as a consistency check, we verified that no cells retained three or more UVIs. Residual conflicts in few cells with exactly two remaining UVIs were resolved conservatively: if both barcodes were detected in other cells, both were discarded; if only one barcode was shared across cells, the private barcode was discarded. Thus, only the interpretable shared barcode was retained when a unique resolution was possible (the other infection may have been true but, occurring once, would not yield a network). The resulting, final F5 assignment served as the final per-cell network barcode used for downstream analyses.

#### Identification of AAV-positive cells

AAV-positive cells were identified based on the number of detected AAV helper-derived transcripts. True Negative cells, which had not been exposed to AAV, were used to estimate the ambient background detection rate. We selected the lowest transcript-count threshold at which fewer than 1% of True Negative cells were classified as AAV-positive (2 transcripts at FPR=0.0025), which was applied to all cells.

#### Reconstruction of UVI-defined networks

Cells sharing a final F5-filtered UVI within the same biological sample constituted one reconstructed network. AAV-positive cells within these networks were classified as starters, and AAV-negative cells as connected cells. Networks were categorized as starter-negative, single-starter, or multi-starter. Analyses requiring defined synaptic directionality were restricted to single-starter networks, in which every connected cell was paired with the unique starter carrying the same UVI. These pairs were counted by starter and connected cell state to quantify presynaptic input composition, with homotypic connections defined as pairs sharing the same fine cell-state annotation. Starter-negative fragments were retained for analyses that did not require identification of the postsynaptic starter (e.g. clone2vec).

#### Organoid-to-organoid reproducibility quantification

Organoid-to-organoid reproducibility was quantified from cell-type composition profiles of rabies-traced cells and matched cell type composition controls. Cells were counted by cell type within each sample or genotype and converted to relative abundances. A small regularization constant (ε:0.01) was added to all categories before renormalization and centered log-ratio transformation. Pairwise similarity was then calculated as weighted Pearson correlation in CLR space, using cell-type weights derived from the geometric mean abundance of each cell type across samples. Correlation matrices were generated separately for traced and control cells at sample and genotype level, and off-diagonal upper-triangle coefficients were used for group-wise comparison by two-sided Wilcoxon rank-sum test.

### Network Analysis

#### Comparison to published transsynaptic tracing studies

Comparative recovery was calculated using two reported quantities across studies: postsynaptic neurons with recovered presynaptic networks and presynaptic neurons assigned to those networks. Published values were extracted from ^34^ Figure 4 and Figure 4B source data; ^35^, main text; and ^36^ Figure 5F; COSCO values correspond to Figure 3G. Fold changes were computed separately for postsynaptic and presynaptic recovery as COSCO/published count and reported as the rounded range across studies.

#### Compositional analysis of cell states in background tissue vs. networks

We quantified the relationship between the proportional abundance of each neuronal cell type in the tissue background (CTC) and among rabies-traced neurons (Traced), across organoids. Analyses included wild-type organoids with neuronal cell types present in at least 500 cells in both CTC and RV^+^ populations. For each sample, we computed fractions of each neuronal cell type within CTC and RV^+^ populations. We then calculated correspondence between tissue and network abundance across samples using two statistics: (i) the slope of a linear model (Network_fraction ~ 0 + Tissue_fraction), measuring how strongly network participation scales with tissue prevalence, and (ii) Spearman correlation coefficient, measuring consistency of network participation across organoids. Slope parameters and correlation coefficients calculated on individual cell types were compiled into a summary scatterplot.

#### Quantification of input diversity and homophily

For each comparison, cell-state composition was represented as a count vector over the same predefined set of 17 annotated cell states, retaining zero-count categories. We quantified compositional inequality using the Gini coefficient and used its complement, 1-Gini, as an input-diversity score. This metric captures how evenly cells are distributed across the fixed cell-state universe, with higher values indicating broader and more even representation. We used 1-Gini because it is bounded, intuitive, and directly interpretable as inverse compositional inequality (Figure 3I). Other diversity metrics, including entropy- or Simpson-based measures, are powerful for comparisons across variable category spaces, or more complex scenarios, but are less straightforward to interpret and/or are unnecessary for our fixed-universe analyses.

#### Comparison of excitatory input architecture to mouse visual cortex

Layer-specific input architecture was compared between day-150 wild-type human cerebral organoids and adult mouse visual cortex using four matched excitatory neuron classes: ExN L2/3, ExN L4, ExN L5, and ExN L6. Organoid analyses were restricted to UVI-defined wild-type networks containing exactly one annotated starter cell. For each network, connected cells were paired with the single starter cell sharing the same UVI, and starter–connected pairs involving the four matched excitatory classes were counted. Organoid ExN L2–4a and ExN L2–4b states were excluded from the cross-system comparison because they lacked direct mouse annotation matches, but were retained in the organoid-only supplementary scatter plot.

Adult mouse visual cortex data ^69^ were reanalyzed after restricting the dataset to glutamatergic neurons from layer-targeted rabies tracing experiments. Injection identities were mapped to L2/3, L4, L5, or L6 starter classes; L5-targeting experiments were collapsed into one L5 group, and L6b cells were merged with L6. Because the mouse dataset was not virally barcoded, all rabies-labeled glutamatergic neurons from each layer-targeted experiment were treated as inputs to the corresponding starter layer.

For each system, starter-by-input count matrices were normalized within each starter class to obtain fractional input composition (per column) across the same four excitatory classes. Homotypic connectivity was defined as the diagonal fraction of this normalized matrix, corresponding to inputs from the same excitatory class as the starter population. Input inequality was quantified as the Gini coefficient of the incoming four-class input distribution per starter class, and input diversity was defined as 1 minus the Gini coefficient. For the organoid-to-mouse overlay, homotypic connectivity and Gini coefficient were z-scored within each dataset across the four matched classes. Raw values were shown (**Figure S3K–N**), and z-scored values were used in Figure 3I.

#### Tenm3 Adgrl2 homotypic attraction

We identified all two-cell networks with exactly one AAV-positive starter and one connected partner. For each pair, we extracted normalized TENM3 and ADGRL2 expression values. Four cross-cell correlation tests (2 homotypic, 2 heterophilic) were computed using Spearman correlation. To assess significance, we generated a 10,000-permutation null by shuffling starter identities while keeping partner expression fixed. Empirical p-values were defined as the fraction of permuted correlations exceeding the observed correlation. We compared observed correlations to their null distributions to determine whether network-specific structure was present. The p-values from the correlation test (cor.test, Spearman) confirmed the results (not shown).

#### Network embedding and motifs

For cluster-free identification of connectivity patterns, we applied clone2vec (c2v) ^38^, an algorithm adapted from the word2vec Skip-Gram architecture. word2vec ^74^ learns a *word’s overall meaning* by taking all sentences with an *occurrence* of this word, and then transforming the *surrounding words’ frequencies* into an embedding vector. Analogously, clone2vec learns each *network’s “meaning”* (its similarities or distances to other networks), by taking the transcriptional kNN-neighborhoods of all *occurrences* of this network (that is each cell in this network), and then aggregating the surrounding network-occurrence frequencies (that is the ***k*** closest cells belonging to any network) into an embedding vector. The clone2vec algorithm was initially developed for the analysis of clones obtained from viral barcoding experiments, but mathematically both synaptic networks and clonal groups can be treated as sets of corresponding cells. The clone2vec workflow comprised the following steps: **Step 1: Cellular Space & Neighborhoods**. We embedded the single-cell expression matrix into a reduced latent space via PCA, followed by batch correction of the space with Harmony ^51^. We then selected cells belonging to networks of size >=3, and constructed a k-Nearest Neighbors (kNN_15_) graph to define a local transcriptomic neighborhood or “context” for these UVI-bearing cells. **Step 2: Generating the Training Corpus**. We then superimposed Unique Viral Identifiers (UVIs) onto each cell in this kNN-graph, replacing individual cell identities with their synaptic network identities. For each cell of the target network, this mapped the target cell to a distribution of its ***k***=15 co-occurring context networks, repeating this for all cells in a network. Repeating this procedure for all networks generated a ***c***^***\****^***N*** (rows:cells ^*^ columns:networks) count matrix, where each row consists of non-negative integers that sum up to ***k***. (Note that for visualization, this is simplified into a ***k***^***\****^***N*** matrix in Fig.4A, and that, for computational efficiency, it is actually stored as an ***N***^***\****^***N*** matrix). **Step 3: Skip-Gram Optimization**. Rather than fitting each target+context vector (rows of ***c***^***\****^***N***) as a single observation, clone2vec decomposed each into **k** individual target-context pairs. The resulting pool of pairs was shuffled across all cells and processed in randomized mini-batches (batch size = 64) for a two-layer Skip-Gram neural network with a latent dimension of ***h*** = 10. This way, during Adam optimization (learning rate = 0.001, full SoftMax output and negative log-likelihood loss, 125 epochs, random seed = 4, single training run), the fitted probabilities reflected the empirical frequency of network co-occurrence. This step is mathematically equivalent to both Skip-Gram and multinomial GLM-PCA ^105^, and can be understood as a low-rank decomposition that preserves maximal variance of the original matrix while accounting for the count nature of its entries. In essence, the neural network uses the target network identity to predict its corresponding context network distribution, and optimization iteratively adjusts its weights via a stochastic gradient descent method to maximize the probability of predicting true co-occurrences. **Step 4: Extracting the Latent Embedding**. After convergence, the model’s input-to-hidden ***N***^***\****^***h*** matrix acts as the low-dimensional latent embedding of each network and can be used for UMAP visualization (n_neighbors = 15, min_dist = 0.2) and Leiden clustering (resolution = 0.75) (similarly to how these are calculated on the primary PCA embedding in standard single-cell analysis).

#### Cell state diversity across network motifs

Cell state diversity within network motifs was quantified separately for starter and connected cells after restricting the analysis to motif-assigned cells from single-starter networks. Starter cells were defined as AAV-positive traced cells and connected cells as AAV-negative traced cells. For each motif group, diversity was calculated across all cell types in the dataset, as 1 minus the Gini coefficient, using a fixed universe of 17 cell types to preserve zero-count categories and ensure comparability across subsets. The identical computational framework was applied to the wild-type and TSC-focused panels to enable direct comparison of motif-resolved diversity patterns across datasets.

#### Motif-specific enrichment of connected cell states

For motif-specific enrichment analyses, we used wild-type DIV120–DIV150, motif-assigned, single-starter networks. Connected cells were defined as rabies-traced AAV-negative cells. Connected cell states represented by fewer than 100 cells were excluded.

For Fig. 4L, connected cells were tabulated by cell state and motif. Expected counts were estimated from the row and column marginals under an independence model, using the connected-cell distribution within the analyzed networks as the background. Motif-specific deviations were shown as Pearson residuals, calculated as observed minus expected counts divided by the square root of expected counts. Tiles with both observed and expected counts below 10 were masked, residuals were clipped at ±30 for visualization, and numeric labels were shown for absolute residuals greater than 2.

For Fig. S4H, the same framework was applied to motif-level all-starter columns and to motif-by-starter-cell-state columns. Starter-resolved columns with fewer than 50 connected cells were excluded. Enrichment was shown as log2 observed/expected counts with a pseudocount of 0.1. Log2 enrichment was chosen as this effect-size measure is comparable across per-motif and per-motif-and-starter-cell-type categories with differing sample sizes, enabling consistent visualization of relative connectivity patterns. All-starter motif columns were masked below 10 observed and expected cells; starter-resolved columns were masked below 5 observed and expected cells. Tile-wise significance was assessed by two-sided Fisher’s exact test followed by FDR correction and annotated at adjusted P < 0.05, P < 0.01, and P < 0.001.

#### Motif assignment validation

To test whether clone2vec motifs could arise from stochastic barcode assignment, we generated three randomized control datasets by independently shuffling UVI labels across UVI-bearing cells within each organoid. This preserved composition, barcode frequency, and network-size distributions, while disrupting the relationship between network identity and transcriptomic neighborhood. Experimental and randomized networks were then jointly processed through the same clone2vec workflow, embedded in network-level clone2vec space, and assigned to motifs by Leiden clustering at resolution 1.5. For each dataset group (experimental, simulation 1, simulation 2, and simulation 3), we counted the number of networks assigned to each motif and compared motif distributions between all dataset pairs using Pearson’s chi-squared test. Pairwise p-values were visualized as –log10(p-values) (Figure S4D). The analysis was then repeated by shuffling across all cells in traced samples (Trace + CTC), which supported the same conclusion.

#### Connectivity-defined transcriptional programs in L4 excitatory neurons

To test whether synaptic network context is associated with transcriptional variation within a single neuronal class, we analyzed L4 excitatory neurons from wild-type DIV150 organoids with clone2vec motif annotations. We first analyzed AAV^+^RV^+^ starter L4 ExNs, and then AAV^-^RV^+^ connected L4 ExNs. Starter-cell analysis was restricted to single-starter networks, linking L4 starter transcription to motif-resolved presynaptic network composition. Connected-cell analysis included all motif-assigned L4 connected cells from networks with zero or one detected starter cell, retaining starter-negative network fragments while excluding multi-starter networks.

For L4 starter cells, motif-wise one-versus-all differential expression was performed across motifs represented by more than 20 L4 starter cells. Differential expression used log-normalized RNA counts and Seurat Wilcoxon tests with min.pct = 0.5, log2 fold-change threshold 1, and Benjamini–Hochberg adjusted p < 0.01. Testing was restricted to expressed coding genes. Significant positive markers were summarized by gene-wise z-scored average expression across motifs and by dot plots showing expression prevalence and scaled expression.

For L4 connected cells, motif-wise one-versus-all differential expression was performed using the same framework, with log2 fold-change threshold 0.5 and Benjamini– Hochberg adjusted p < 0.01. Motif-level marker expression was hierarchically clustered across motifs, defining a Deep Motif Group comprising m0, m1, and m4, and an Upper Motif Group comprising m2, m3, m5, m6, and m7. These groups were then compared directly by pairwise differential expression on log-normalized RNA counts with min.pct = 0.5, absolute average log2 fold change > 0.5, and adjusted p < 0.01.

Starter-cell marker interpretation used selected GO and SynGO annotations related to synapse organization, cell adhesion, axonal, dendritic, presynaptic, and postsynaptic compartments. For connected-cell motif groups, Deep- and Upper-associated genes were analyzed separately for GO Biological Process enrichment using clusterProfiler::enrichGO with org.Hs.eg.db, Entrez identifiers, and Benjamini–Hochberg multiple-testing correction. Enriched terms were simplified by semantic similarity filtering at cut-off 0.7. GO overlap between Deep and Upper Motif Groups was quantified by gene ratio per term, and selected shared terms were resolved at the gene level by plotting the corresponding Deep-versus-Upper differential-expression effect sizes.

#### Transcriptomic subclustering of L4 excitatory neurons and connectivity enrichment

To test whether connectivity-associated neuronal states could be recovered from transcriptomes alone, we independently reprocessed all L4 excitatory neurons from DIV150 H9, 177, and TSC2 isogenic control organoids using the standard Seurat workflow described above. The clustering solution at resolution 0.2 was used for visualization, resolving three major subclusters and a small additional population. Motif annotations were not used for clustering and were overlaid only after the transcriptomic subclusters had been defined. We used the above motif-group-assignments and Deep Motif Group membership was projected onto the L4 UMAP by weighted kernel density estimation using Nebulosa with an adjustment factor of 2. For each major transcriptomic subcluster, the fraction of cells belonging to each motif group was normalized to the corresponding fraction across all motif-assigned L4 neurons and expressed as a percentage of the dataset-wide expectation, with 100% indicating no enrichment.

#### Analysis of TSC and isogenic control organoids

##### Cell state and motif fractions

To quantify lineage and network composition changes in TSC, we analyzed the data subset corresponding to **Figure 6**, comprising TSC2-mutant organoids and matched isogenic controls. Within each organoid sample, cell-state fractions were calculated for both CTC and Traced populations. Control and mutant samples were compared using a two-sided median permutation test with 100,000 permutations, and p-values were adjusted across tested cell states using Benjamini–Hochberg correction. The same per-sample framework was applied to motif composition in Traced cells. For the aggregate upper-layer analysis, ExN L4, ExN L2–4a, and ExN L2–4b fractions were summed per Traced sample and compared between groups using the same permutation test.

##### Homotypic connectivity and network-level diversity in TSC

Homotypic connectivity was quantified in individual UVI-defined rabies-traced networks containing exactly one AAV-positive starter cell and at least one connected cell. Each valid network was decomposed into all starter-connected cell pairs, and each pair was classified as homotypic when the starter and connected cell shared the same fine cell-type annotation. A two-sided Fisher’s exact test was performed, and the average for each condition was displayed.

Individual-network diversity was quantified in single-starter networks containing at least five cells. For each network, cell-type composition was calculated across the full cell-type annotation set. Diversity was then quantified as 1 minus the Gini coefficient over all 17 cell types in the dataset and analyzed separately for networks of L4 ExN starters or any starters, and Control and TSC distributions were compared using a two-sided Wilcoxon rank-sum test (no significant difference found).

## Supplementary Figures

**Fig. S1.**
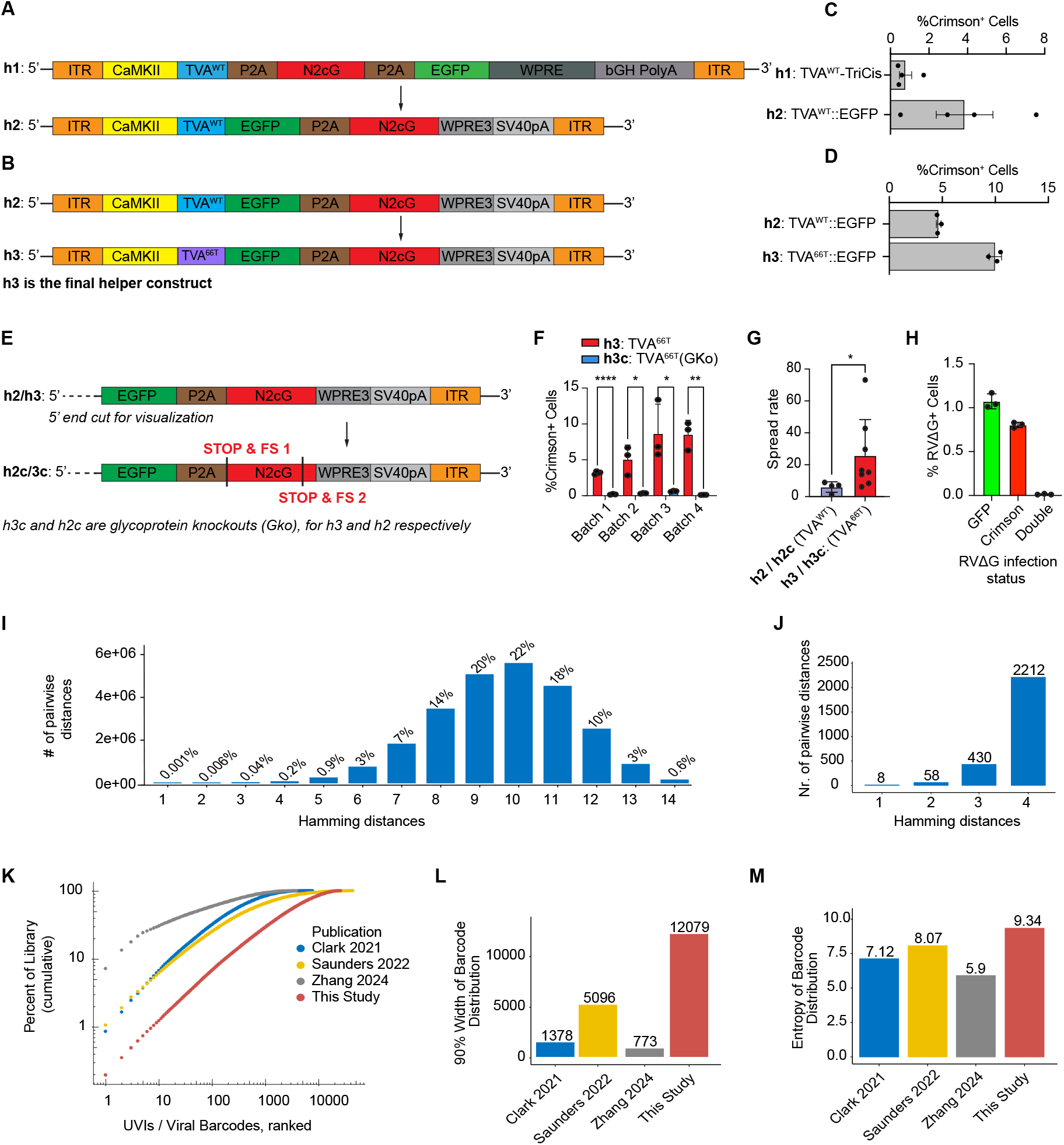
Optimization of starter constructs, barcode diversity, and controls for tracing. **(A)** Schematics of the tricistronic (TVA^WT^-TriCis) and the optimized, bicistronic, TVA^WT^::EGFP fusion starter constructs tested in panel C. The replacement of the bovine growth hormone polyadenylation signal and the full WPRE regulator shortened the construct length from AAV’s packaging limit 4.9 kb to 4.0 kb. **(B)** Schematics of TVA^WT^::EGFP and TVA^66T^::EGFP starter constructs tested in panel D. **(C)** TVA::EGFP fusion improves transsynaptic spread efficiency. FACS-based quantification of the proportion of E2-Crimson^+^ cells comparing TVA^WT^-TriCis and TVA^WT^::EGFP constructs (N=2, n=4 each; mean ± SEM). **(D)** TVA^66T^ increases transsynaptic spread relative to TVA^WT^. FACS-based quantification of the proportion of E2-Crimson^+^ cells across constructs (N=1, n=3 each; mean ± SD). **(E)** Schematics of TVA^66T^::EGFP and TVA^66T^(GKo) AAVrg constructs. In TVA^66T^(GKo), dual frameshift mutations introduce premature stop codons in N2cG, abolishing glycoprotein complementation and prevent transsynaptic spread. **(F)** Proportion of E2-Crimson^+^ cells comparing TVA^66T^ (n=13) and TVA^66T^(GKo) (n=12) across four independent organoid differentiation experiments (N=4; DIV 120, mean ± SD; ^*^p < 0.05, ^**^p < 0.01, ^****^p < 0.0001). **(G)** Spread rate defined as the ratio of % E2-Crimson^+^ cells in glycoprotein-complemented versus GKo conditions, comparing TVA^WT^ (n=4) and TVA^66T^ (n=8) constructs. (Mean ± SD. ^*^p < 0.05, Wilcoxon rank-sum test) **(H)** FACS quantification of dual-color rabies infection assay showing RVΔG-EGFP^+^, RVΔG-E2-Crimson^+^, and double-positive cells after co-infection by equal viral dosages (1^*^2e^5^ or 2^*^1e^5^ virions/well each) corresponding to twice the viral dose used for tracing (n = 3; mean *±* SD). **(I)** Pairwise Hamming distance distribution across a random subset of 5,000 sequence-corrected UVIs in the RVΔG–UVI–E2-Crimson library. Percentages indicate the proportion among all barcode pairs. High sequence divergence shields against PCR or sequencing error-driven UVI misassignment. **(J)** Absolute counts of pairwise barcode distances at the four lowest pairwise Hamming distances (zoom-in of I) **(K)** Cumulative rank-frequency distribution of UVIs supported by >2 reads from the RVΔG–UVI library versus published datasets. **(L, M)** 90% library width (q90WL), and Shannon entropy of barcode distributions across studies (Methods). q90WL is defined as the minimum number of UVIs required to account for 90% of reads in any sample (Methods). N=number of independent differentiation experiments. n=number of organoids or independent replicates as indicated.

**Fig. S2.**
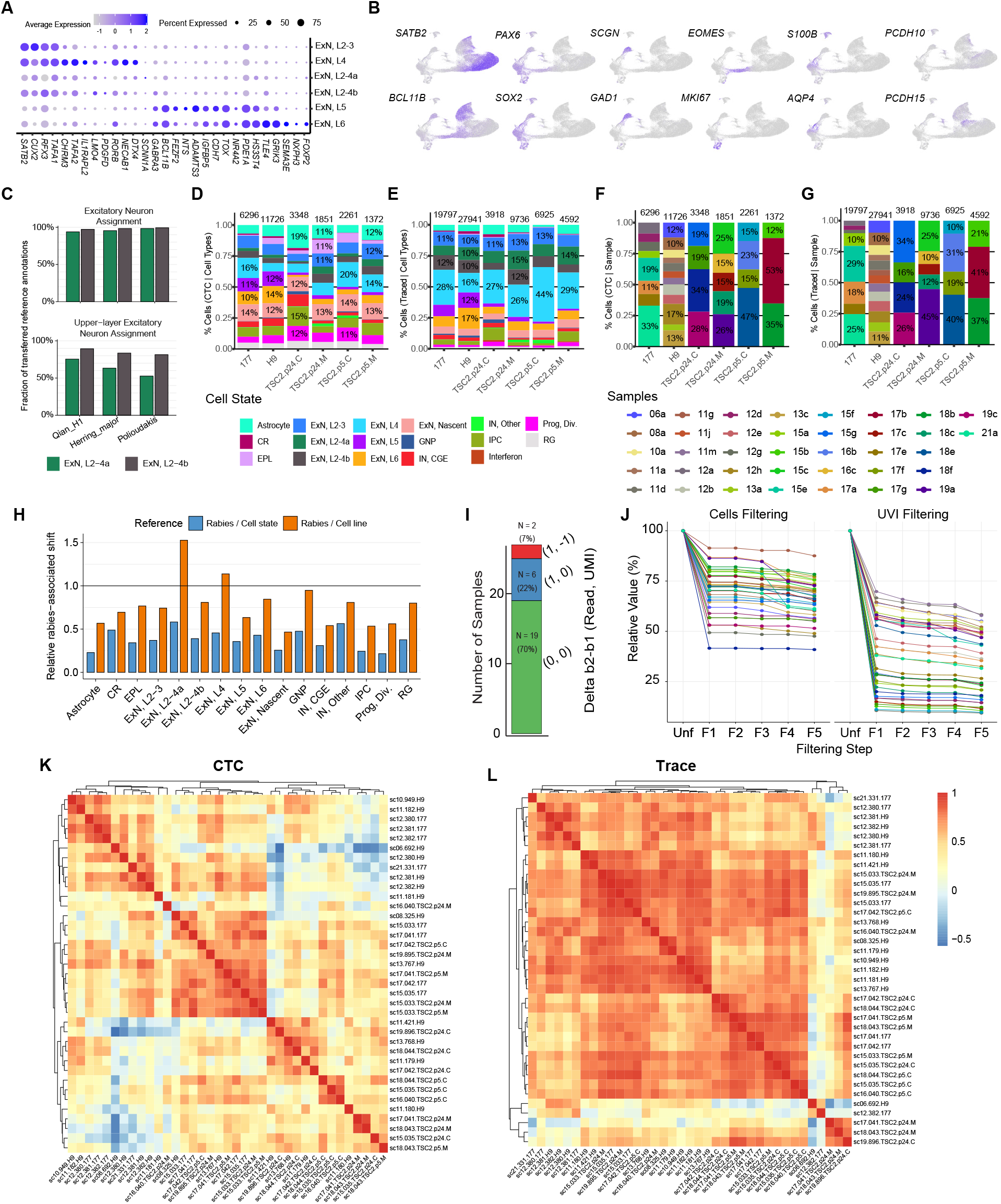
Cell-state annotation and compositional validation of COSCO-traced populations. **(A)** Dot plot of marker genes confirming excitatory neuron lineage annotations. Fill indicates log-scaled expression; size indicates percentage of expressing cells. **(B)** Selected cell-state marker expression on UMAPs. **(C)** Fractions of matching excitatory-neuronal, or upper-layer-excitatory-neuronal annotations transferred onto cell states ExN, L2-4a and b across three fetal-brain reference datasets. **(D, E)** Cell-state proportions across genotypes in CTC (D) and Traced (E) populations. **(F, G)** Cell-state proportions across samples in CTC (F) and Traced (G) populations. **(H)** Rabies-associated root-mean-square (RMS) log2 fold changes in gene expression Trace and CTC cells, normalized either to RMS log2 fold change between cell states (blue) or to change between cell lines within a cell state (orange). Values below 1 indicate the rabies-associated shift is smaller than the reference change. **(I)** Comparison of thresholds resulting from UVI-threshold optimization by the A-B1 or A-B2 strategies. Bars show Δ(B2–B1) for the optimized (R, U) threshold pairs. B1 and B2 selected identical thresholds in 19/27 libraries (70%). B2 selected a one-step higher read threshold in 6/27 libraries (22%); and it resulted in a 1:1 R–U trade-off in 2/27 libraries (7%). Thus, B2 selected equivalent or more stringent read-depth thresholds in all libraries and was used for final library-specific UVI calling. **(J)** UVI and cell dropout per filtering step across scRNA-seq samples; lines colored by samples. **(K, L)** Pearson correlation coefficients of CLR-transformed cell type proportions across individual samples in CTC (K) and Traced (L) cells, as in Figure 2K-L. Heatmaps share a common color scale.

**Fig. S3.**
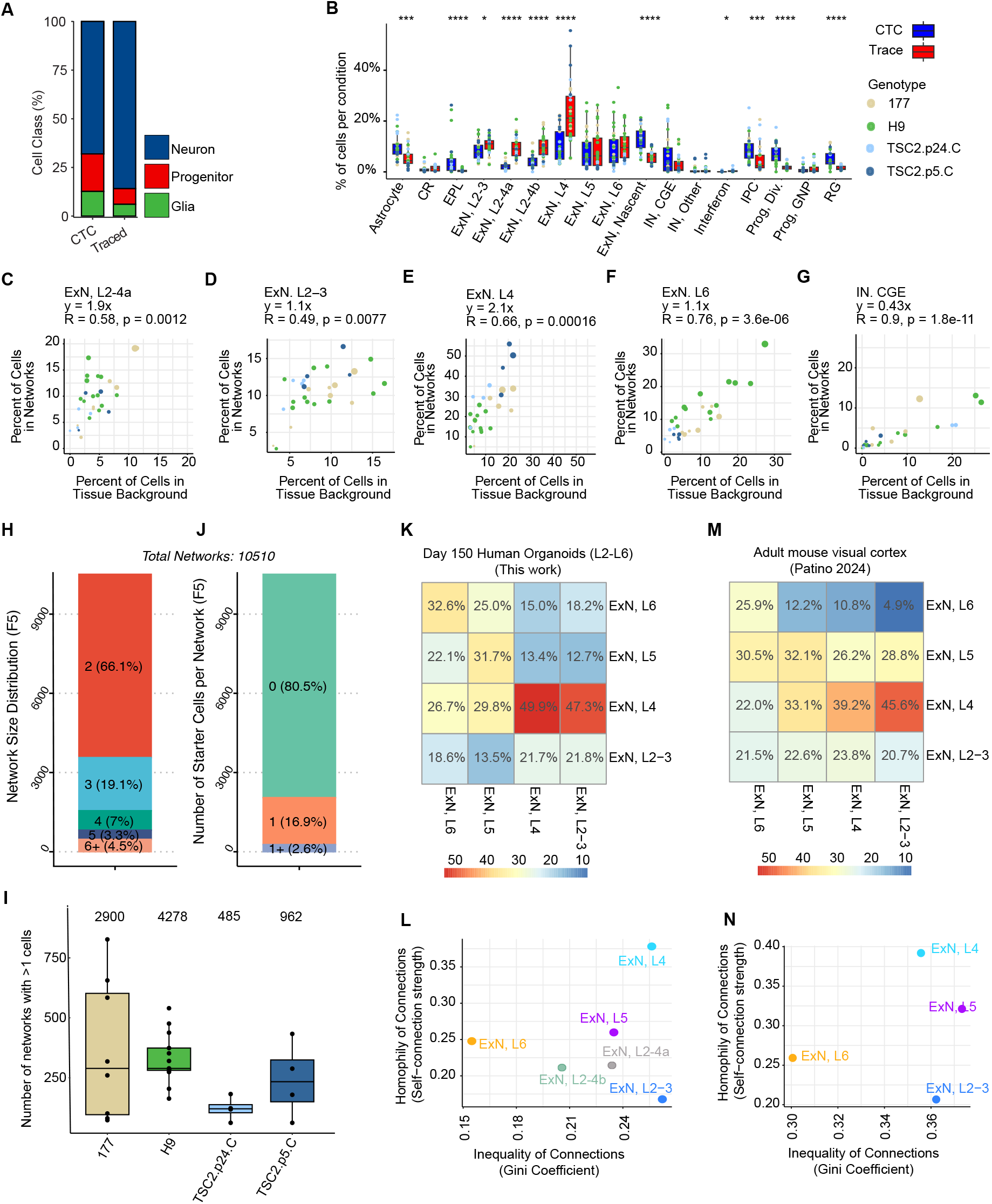
Cell-type-specific connectivity patterns, organoid-to-mouse conservation, and network structure in organoids. **(A)** Cell class composition in CTC and Traced compartments. **(B)** Percentage of cells per CTC and Traced conditions across cell states in healthy controls. Two-sided Wilcoxon rank-sum test with Benjamini-Hochberg adjusted p-values. ^*^p < 0.05, ^***^p < 0.001, ^****^p < 0.0001. H9: N=5, n=13; 177: N=4, n=8; TSC2.p24.C: N=3, n=4; TSC2.p5.C: N=2, n=4. **(C–G)** Relationship between cell type abundance in Traced (network) over CTC (tissue) compartments for additional neuronal subtypes. Each point represents a sample, colored by genotype (B). **(H–J)** Characteristics of the final set of 10,510 networks. (H) Distribution of network sizes. (I) Number of networks per sample (dots) across four healthy control lines. Numbers above each box indicate the total number of networks per cell line. (J) Distribution of the number of starter cells detected per network. Sample sizes are as in (A). **(K)** Presynaptic input fractions (rows) per starter types (columns) across canonical cortical excitatory neurons, in cerebral organoids. **(L)** Homotypic connectivity strength versus input inequality (Gini coefficient) in organoids. Cell states without matching annotation in the mouse cortex (ExN L2–4a and ExN L2–4b) shown as opaque points. **(M–N)** As in K–L, but from the adult mouse visual cortex ^69^. N = number of independent differentiation experiments. n = number of scRNA-seq samples.

**Fig. S4.**
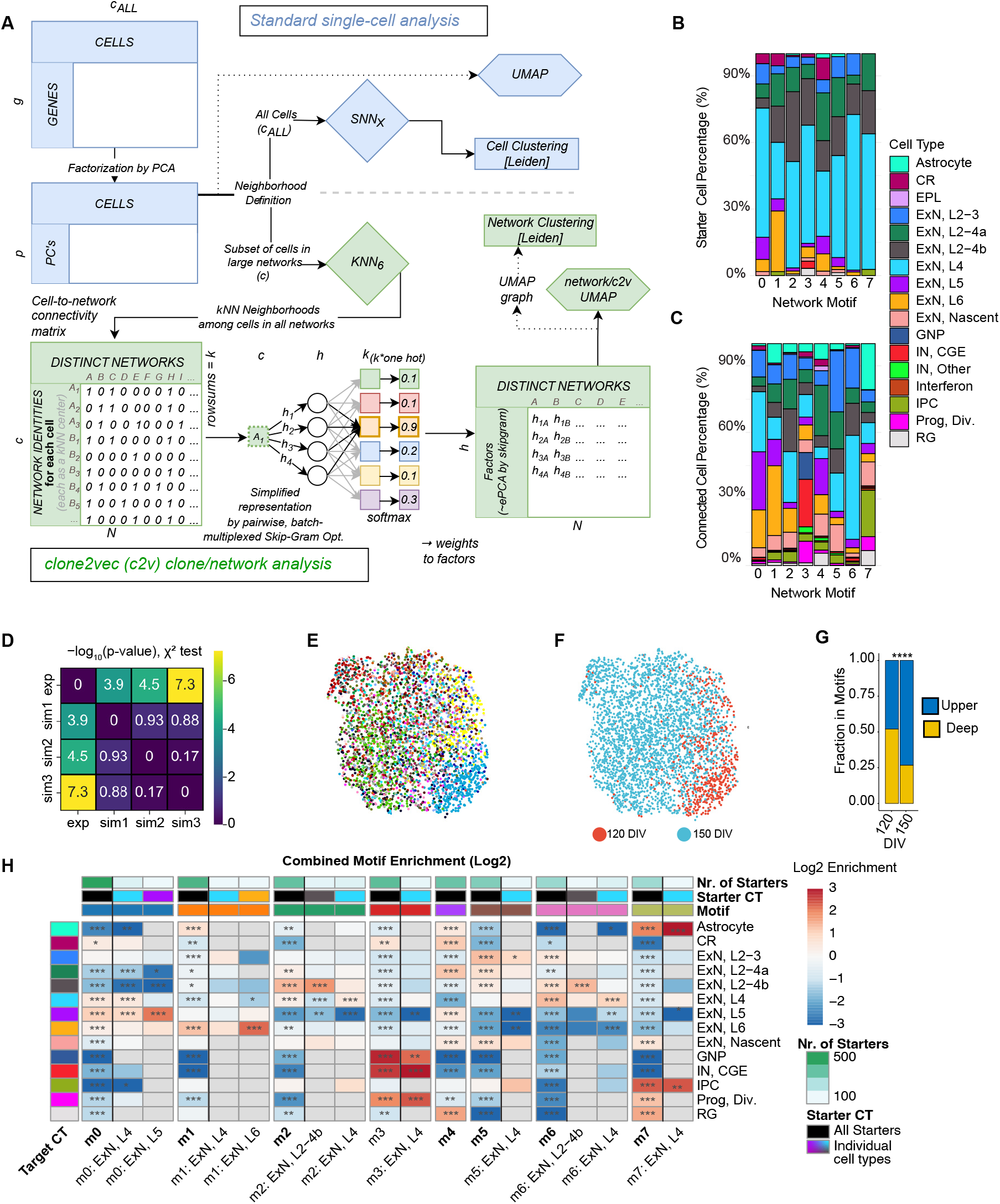
Computational validation and compositional characterization of clone2vec network motifs. **(A)** Step-by-step schematic of c2v data transformations. Transcriptomic similarity is quantified in PCA space, followed by kNN-based neighborhood definition. Network identities are assigned to each cell, transforming transcriptional similarity into a structured 1+k training set of co-occurring network IDs. A Skip-Gram model with one-hot encoded, batch-based Adam optimization with SoftMax learns factorized weights representing each network’s cellular context. This produces a dense latent embedding in which networks with similar cellular compositions cluster together. **(B, C)** Distribution of starter (B) and connected (C) cells across network motifs and cell states. **(D)** Chi-squared test with three shuffled-barcode controls for clone2vec motif assignments indicate that observed motif structure is not explained by chance. Heatmap values show –log10(p-values) from pairwise chi-squared tests comparing motif distributions between dataset groups. Randomized controls were mutually similar, and all significantly differed from experimental networks. **(E)** Distribution of samples across network motifs. H9: N=5, n=13; 177: N=4, n=8; TSC2.p24.C: N=3, n=4; TSC2.p24.M: N=3, n=5; TSC2.p5.C: N=2, n=4; TSC2.p5.M: N=2, n=3. **(F)** Distribution of organoid age across network motifs (Turquoise: 150 DIV: N=17, n=35; Red: 120 DIV: N=2, n=2). 120 DIV samples correspond to the yellow and blue sample labels in panel E. **(G)** Fraction of networks per motif across organoid ages, grouped by deep (L5–6 ExNs) or upper layer (L2–L4 ExN) neuronal composition. **(H)** Combined heatmap of log2 observed/expected enrichment for connected cell states across motif-level all-starter columns and motif-by-starter-cell-state columns. Expected counts were calculated from the corresponding independence model. Starter-resolved columns with fewer than 50 connected target cells were excluded. Top bar annotations indicate motif identity, starter-cell type, and the number of single-starter networks contributing to each column. Asterisks indicate two-sided Fisher’s exact tests with FDR correction: ^*^ P_adj_ < 0.05, ^**^ P_adj_ < 0.01, ^***^ P_adj_ < 0.001. Gray tiles indicate low-count entries where both observed and expected counts were below 5. N = number of independent differentiation experiments. n = number of scRNA-seq samples.

**Fig. S5.**
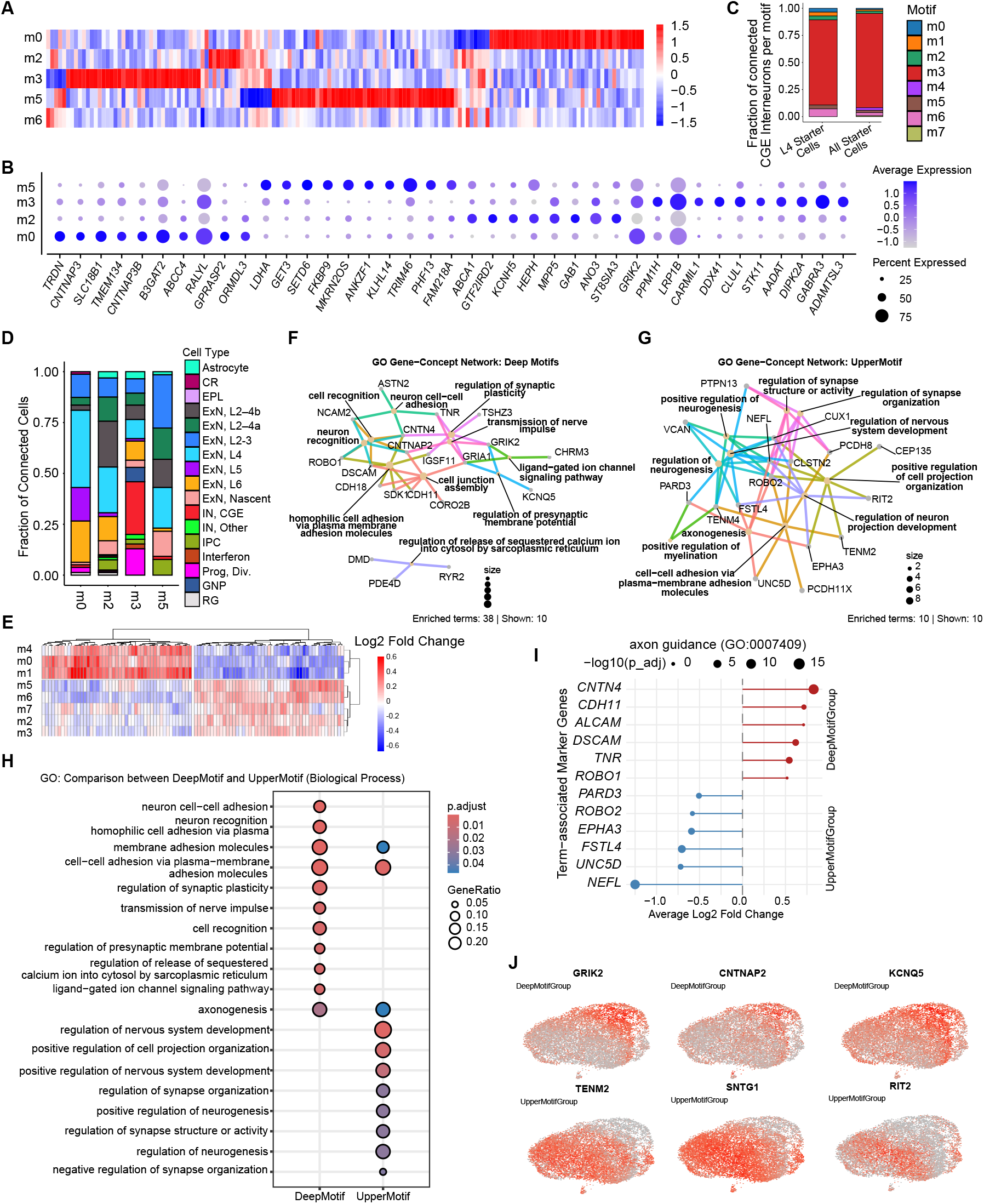
Transcriptional specialization of connectivity motifs in L4 excitatory neurons. **(A)** Heatmap of motif-specific marker genes in L4 starter cells, shown as gene-wise z-scored average expression across motifs. **(B)** Dot plot of top differentially expressed genes in L4 starter cells across motifs with sufficient starter cell numbers (>20 L4 starters), corresponding to Figure 5C. Point size indicates percentage of expressing cells; color indicates average scaled expression. **(C)** Distribution of connected CGE interneurons across motifs for networks with L4 ExN starters or with any starter cell. In both groups, most CGE interneuron inputs occur in motif 3 (m3). **(D)** Cell-state composition of presynaptic partners of L4 starter cells across the motifs used for starter-cell DGEA, showing motif-specific differences in input cell-state abundance. **(E)** Heatmap of genes identified by pairwise Deep versus Upper Motif Group DGEA in L4 connected cells, displayed as average scaled expression across individual motifs. Genes were selected by absolute log2FC > 0.5 and adjusted p < 0.01. This confirms that the group-associated transcriptional signatures are shared across nearly all constituent motifs. (**F, G**) GO Gene-Concept networks for the Deep Motif Group (F) and Upper Motif Group (G). Nodes represent enriched GO terms or contributing genes; edges indicate gene-to-term membership. Deep Motif Group genes are enriched for cell recognition and synaptic plasticity; Upper Motif Group genes for axonogenesis and nervous system regulatory programs. (**H**) Dot plot of GO Biological Process enrichment overlap between Deep and Upper Motif Groups. Point size indicates gene ratio, color indicates adjusted p-value. **(I)** Lollipop plot of average log2 fold change for “axonogenesis” (GO:0007409) marker genes distinctly utilized by Deep or Upper Motif Groups. Point size indicates *−*log10 adjusted DGE p-value, direction indicates Deep versus Upper Motif Group association. **(J)** UMAP plots of representative Deep Motif Group markers (GRIK2, CNTNAP2, KCNQ5) and Upper Motif Group markers (TENM2, SNTG1, RIT2) across L4 neurons. Red intensity indicates normalized expression on the reprocessed L4 excitatory neuron UMAP. N = number of independent differentiation experiments. n = number of scRNA-seq samples. p-values in (F–G, H–I) were corrected by the Benjamini-Hochberg method.

**Fig. S6.**
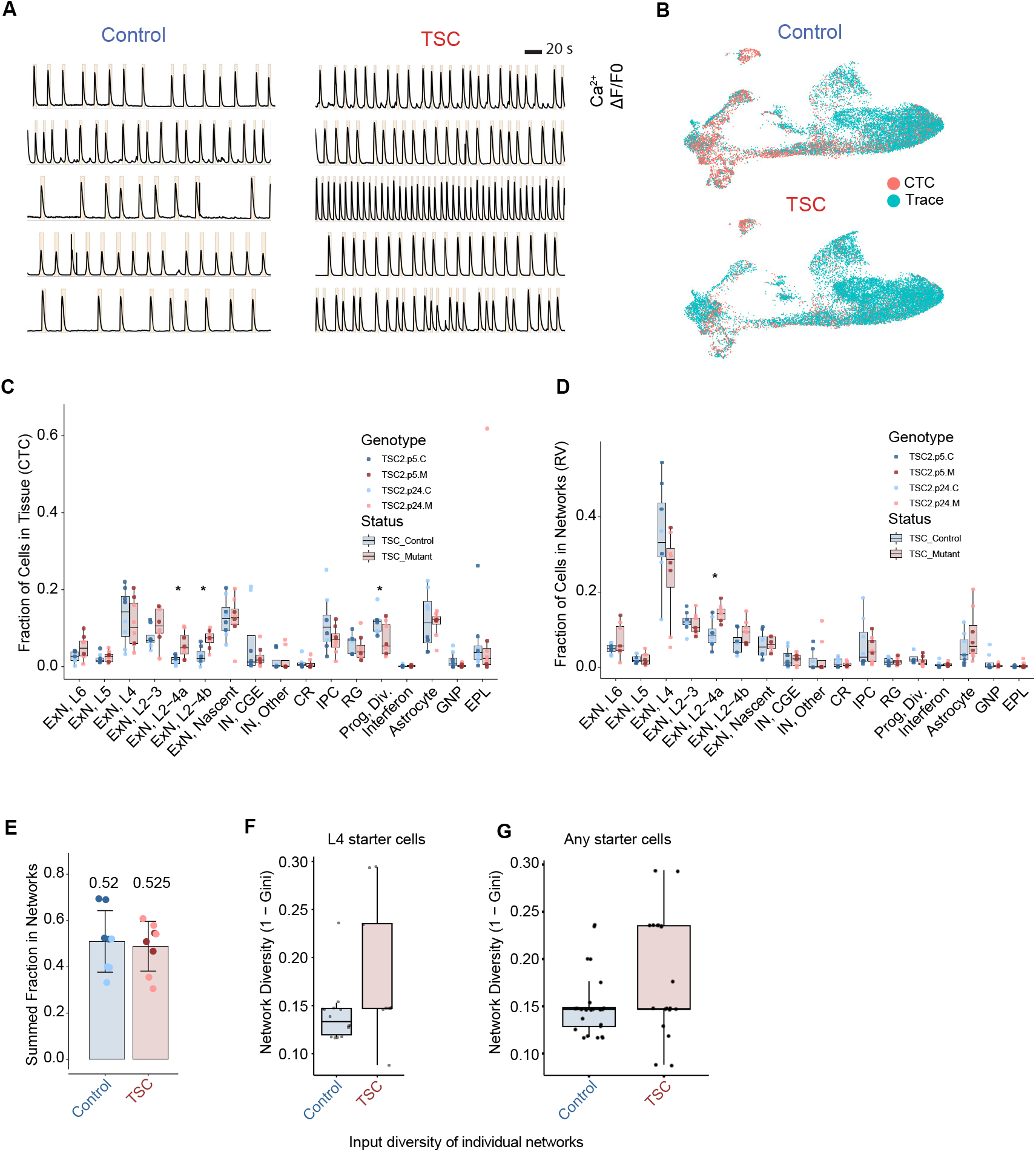
Functional and structural characterization of aberrant connectivity in TSC organoids. **(A)** Additional GCaMP6s activity traces corresponding to Figure 6A. Traces are from separate organoids (one FOV), illustrating spontaneous synchronous network event differences between Control and TSC. Yellow boxes highlight detected network events. **(B)** UMAPs depicting Control (top) and TSC (bottom) cells separated by CTC (red) and Trace (blue) conditions. TSC2.p24.C: N = 3, n = 4; TSC2.p24.M: N = 3, n = 5; TSC2.p5.C: N = 2, n = 4; TSC2.p5.M: N = 2, n = 3. **(C, D)** Fractions of individual cell states in the CTC (C) and rabies-traced (D) populations of TSC and control organoids. Points represent individual samples and are colored by genotype. ^*^FDR-adjusted p < 0.05, two-sided median permutation test on per-sample fractions. Samples as in Figure 6D–E. **(E)** Summed fraction of upper-layer excitatory neurons (ExN L4, L2–4a, L2–4b) in the Traced population shows that the expansion of ExN L2–4a and L2–4b offsets the depletion of ExN L4s, consistent with redistribution within the upper-layer excitatory pool. Samples as in Figure 6D–E. Statistical significance was assessed by a two-sided median permutation test on per-sample summed fractions. **(F, G)** Diversity in single-starter networks with *≥*5 cells from L4 ExN starters (F) or all starters (G) in Control and TSC organoids. Higher diversity values indicate more heterogeneous cell-type composition. Dots denote individual networks. Networks from samples as defined in (B). N=number of independent differentiation experiments; scRNA-seq: n=number of scRNA-seq samples. For all box plots: boxes show median and IQR; whiskers extend to 1.5*×* IQR.

